# Leveraging archaeological specimens to investigate failed invasions: A case study on Okhotsk pigs, northern Japan

**DOI:** 10.64898/2026.04.22.720108

**Authors:** Takumi Tsutaya, Taichi Hattori, Rin Onishi, Chelsea E. Budd, Hideki Minoshima, Tomonari Takahashi, Yu Hirasawa, Sayaka Chiku, Takayuki Omori, Kohei Yamazaki, Minoru Yoneda, Daisuke Kubo, Hajime Ishida, Takao Sato, Rick J Schulting, Hirofumi Kato, Andrzej W Weber

## Abstract

Invasive species pose a major threat to biodiversity, yet our understanding of failed invasions/translocations, instances where alien/introduced species fail to establish, remains limited. Investigating the factors behind failed invasions is critical for improving prevention and management strategies for modern biological invasions. Here, we propose a novel framework that utilizes archaeological archives to uncover evidence of failed invasions. We estimated the biological and ecological factors contributing to the failed invasion of pigs from later prehistory to recent times (299 cal BC to 1900 AD) on Rebun Island in far northern Japan by synthesizing the evidence obtained from stable isotopes, zooarchaeology, and historical documents. Despite the anthropogenic introductions of pigs into Rebun Island, pigs did not establish a feral population and disappeared after ca. 1200 AD. We identified reduced propagule pressure, abiotic resistance due to the cold climate, and decreased resources as the three key factors that contributed to the disappearance of pigs. Pigs are one of the most widespread invasive species globally, and this study represents a novel approach to studying failed invasions using archaeological data, which aligns with the framework of conservation paleobiology.

## Introduction

Invasive species are among the most significant global threats to biodiversity, contributing to the extinction of numerous taxa, particularly on islands (Bellard et al., 2016; Doherty et al., 2016; Jones et al., 2016; Spatz et al., 2017). Effective prevention and management of invasive species necessitate a thorough understanding of the interactions between these species and their environments. However, there is a notable gap in our knowledge regarding failed invasions—instances where alien species do not establish themselves as invasive (Zenni and Nuñez, 2013). This gap is largely due to the difficulty in detecting such cases. Understanding the factors leading to failed invasions is crucial, as it could enhance their effective control (Zenni and Nuñez, 2013).

This study presents a novel framework that leverages archaeological archives to provide detailed evidence of failed invasions, offering insights that could be critical in preventing future cases. The use of paleontological, archaeological, or historical archives for biodiversity conservation is encompassed within the field of conservation paleobiology (Barnosky et al., 2017; Boivin and Crowther, 2021). Interdisciplinary investigations of these archives allow for the reconstruction of species extinctions, habitat shifts, and anthropogenic impacts over timescales ranging from decades to millennia—timescales that are challenging to study using contemporary ecological methods (Barnosky et al., 2017; Boivin and Crowther, 2021). Additionally, insights gained under past climatic regimes and human activities that differ from the present offer an advantage in predicting future biodiversity changes due to climate change (Fordham et al., 2020). While invasive species have been studied within the framework of conservation paleobiology (Crees and Turvey, 2015; Dietl et al., 2015; Gillson et al., 2008; Kim et al., 2023), research focusing specifically on failed invasions remains unexplored (but see Duffy et al., 2000).

The introduction and subsequent extirpation of domesticated pigs (*Sus scrofa*) during the Okhotsk period (6th–13th centuries) on Rebun Island (Supplementary Figure S1) in subarctic Japan is the focus of this study. Pigs are one of the most widespread and impactful invasive species globally, ranking among the top five contributors to vertebrate extinctions (Bellard et al., 2016) and the fourth most invasive mammal (Doherty et al., 2016). Their omnivorous diet and destructive rooting behavior significantly disrupt ecosystems (Barrios-Garcia and Ballari, 2012) and threaten 672 taxa across 54 different countries worldwide (Risch et al., 2021).

Rebun Island is a small island off the northwest tip of Hokkaido, Japan, with an area of 81 km^2^, characterized by continuous human occupation from the Jomon period (from ca. 300 BC and earlier) to the historical Ainu period (ca. until 1900 AD) (Table 1; Supplementary Text S1). During the Okhotsk period, domesticated pigs were introduced, but they disappeared from the archaeological record shortly thereafter (Hattori et al., 2021; Hudson, 2004; Nomura and Utagawa, 2003). The archaeological site of Hamanaka 2 provides a continuous record of human occupation from the Jomon to the Ainu periods (Junno et al., 2021). In the initial Okhotsk period (489–573 cal AD), the Okhotsk people, who migrated from the Amur River basin via Sakhalin (Sato et al., 2021), introduced pigs from Eurasia to the island (Watanobe et al., 2011). These pigs, which had been fully domesticated and distinct from wild boars, did not establish a lasting presence (Hattori, 2017), unlike the domesticated dogs, which were consistently present throughout the cultural sequence (Onishi, 2015) (Table 1; Supplementary Text S1).

**Table 1.**
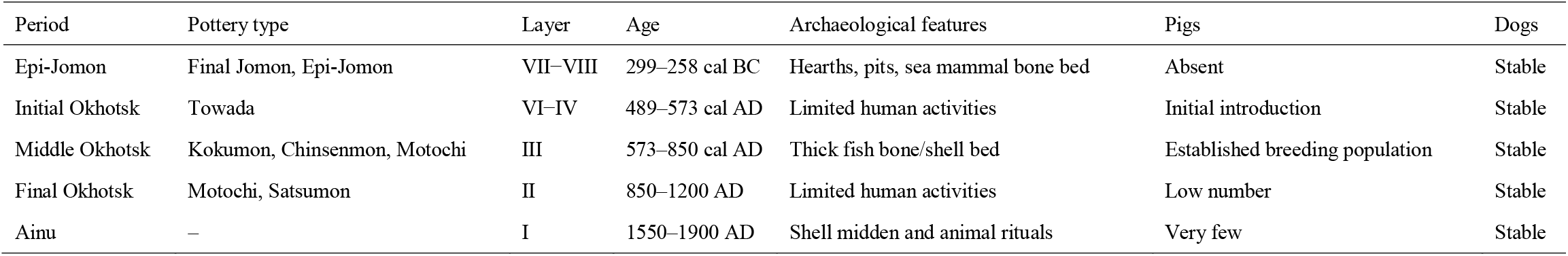
Cultural sequence, their age, and details in the Hamanaka 2 site.

| Period | Pottery type | Layer | Age | Archaeological features | Pigs | Dogs |
| --- | --- | --- | --- | --- | --- | --- |
| Epi-Jomon | Final Jomon, Epi-Jomon | VII–VIII | 299–258 cal BC | Hearths, pits, sea mammal bone bed | Absent | Stable |
| Initial Okhotsk | Towada | VI–IV | 489–573 cal AD | Limited human activities | Initial introduction | Stable |
| Middle Okhotsk | Kokumon, Chinsenmon, Motochi | III | 573–850 cal AD | Thick fish bone/shell bed | Established breeding population | Stable |
| Final Okhotsk | Motochi, Satsumon | II | 850–1200 AD | Limited human activities | Low number | Stable |
| Ainu | – | I | 1550–1900 AD | Shell midden and animal rituals | Very few | Stable |

Previous zooarchaeological studies of Hamanaka 2 have revealed patterns of animal management (Supplementary Text S1). Hamanaka 2 pigs showed evidence of an established breeding population by the early Middle Okhotsk period (573–850 cal AD) and a higher proportion of young individuals (i.e., individuals with unerupted second molars comprised ∼44%: Supplementary Text S1). Pigs also exhibited a greater prevalence of stress markers, such as enamel hypoplasia (EH), particularly after the latter part of the Middle Okhotsk period, compared with wild boars from Jomon archaeological sites on the main island of Honshu (Hattori, 2017). Despite a peak in the number of excavated pig specimens during the Middle Okhotsk period, the species failed to thrive and eventually disappeared. In contrast, dogs showed little stress and no significant morphological changes, and were consistently present throughout the cultural sequences on Rebun Island, indicating a successful adaptation to the island’s environment (Onishi, 2015).

This study aims to uncover the biological and ecological factors contributing to the failed invasion of pigs on ancient Rebun Island using stable isotope analysis. Stable carbon and nitrogen isotope ratios systematically differ among food sources in different categories (e.g., C3 and C4 plants (the latter not present on Rebun), terrestrial animals, and marine fish) and can quantitatively reveal individual differences in diet and habitat use (Makarewicz and Sealy, 2015; West et al., 2006). Stable isotope analyses have been utilized in studies of biological invasion in modern organisms (McCue et al., 2020; Tillberg et al., 2007; Vander Zanden et al., 1999), but few cases have been reported in archaeological contexts (Hofman and Rick, 2017; Swift et al., 2017). By analyzing carbon and nitrogen isotope ratios (δ^13^C and δ^15^N values) of 177 ancient non-human animal bones, including 33 pigs, 13 human skeletons, and modern plant and marine invertebrate tissues, as well as previously reported values of 242 marine fish bones, we investigate dietary changes, habitat use, and interspecies interactions that may have led to the failure of resilience and consequent disappearance of pigs from the island.

## Materials and Methods

The materials and methods are briefly summarized below; the full details can be found in Supplementary Text S4.

Archaeological skeletal materials were obtained through excavation campaigns at Hamanaka 2 from 2011 to 2016 (Kato et al., 2012, 2015; Kato and Iwanami, 2014; Kato and Naganuma, 2016, 2017). Stable isotope ratios of bone collagen were newly obtained from a total of 33 pigs, 83 dogs, 11 humans, and 61 other faunal specimens. Previously reported carbon and nitrogen stable isotopic data on two human (Okamoto et al., 2016; Uchida-Fukuhara et al., 2024) and 242 fish (Tsutaya et al., 2018, 2022) bone samples from Hamanaka 2 were merged into the dataset of this study. Bones excavated from layers VIII–VII, VI–IV, III, and II were assigned to the Epi-Jomon period and to the Initial, Middle, and Final phases of the Okhotsk period, based on the pottery typology and radiocarbon dating (Table 1; Junno et al. 2021). Modern reference samples of plants and marine shellfish were obtained from the summer to autumn of 2016 from Rebun Island. When comparing the modern data with archaeological ones, a +1.0‰ correction for the δ^13^C values was applied to correct the Suess effect (Friedli et al., 1986).

Collagen was extracted from bone samples weighing 6–360 mg using the method described in Tsutaya et al. (2017, 2018). Edible parts were used for stable isotope analysis for modern food samples. Carbon and nitrogen stable isotope ratios were measured using an elemental analyzer–isotope ratio mass spectrometry. Based on repeated measurements of the calibration standards, precision was determined to be less than ±0.1‰ standard deviation (SD) for both δ^13^C and δ^15^N.

A sequential carbon and oxygen stable isotope analysis was conducted on the tooth enamel of pig specimen 2011HA1894, which showed intermediate to slightly elevated δ^15^N values among the isotopic distribution of the Hamanaka 2 pigs. The enamel sampling method followed the protocol described in a previous study (Frémondeau et al., 2012). Carbon and oxygen isotope ratios of enamel were measured by using a Kiel device coupled with an isotope ratio mass spectrometer. The measurement precision (standard deviation) was ±0.01‰ for carbon isotopes and ±0.04‰ for oxygen isotopes.

Radiocarbon ages were measured from approximately 2.5 mg of extracted collagen by using accelerator mass spectrometry. Radiocarbon ages were calibrated with the software OxCal (Bronk Ramsey, 1995) and atmospheric and marine data sets (IntCal20 and Marine20, respectively: Reimer et al., 2020; Heaton et al., 2020). The local marine reservoir effects (⊿R = 217 ± 35), recalculated with Marine20, measured for the Gulf of Aniva, Sakhalin, were used for correction (Yoneda et al., 2007).

The isotopic results were analyzed and visualized by using R software, version 4.2.3 (R Core Team, 2023). Dietary protein contributions from different food categories were calculated using the R package SIAR, version 4.2 (Parnell et al., 2010), with fixed isotopic offset values (Lee-Thorp, 2008).

## Results

### Overall trend

After the removal of poorly preserved specimens based on atomic C/N ratio (DeNiro, 1985) and yield (van Klinken, 1999) of extracted collagen, bone collagen extracted from ancient remains ranged from -21.1‰ to -10.3‰ in δ^13^C and 6.9‰ to 20.8‰ in δ^15^N values (Table 2; Figure 1; Supplementary Tables S1–S4; Supplementary Figures S2 and S3). The isotope ratios for each wild species were consistent with their known diets and ecological niches, aligning with previous studies (Naito et al., 2010; Tsutaya et al., 2014), and confirming the reliability of our results. As expected, marine animals had higher isotope ratios compared to terrestrial animals, with marine mammals showing elevated δ^15^N values due to their higher trophic levels (Figure 1). Modern food samples (i.e., C3 plants and marine invertebrates) showed consistent isotopic values with those expected from their ecological niche (Supplementary Table S5).

**Table 2.**
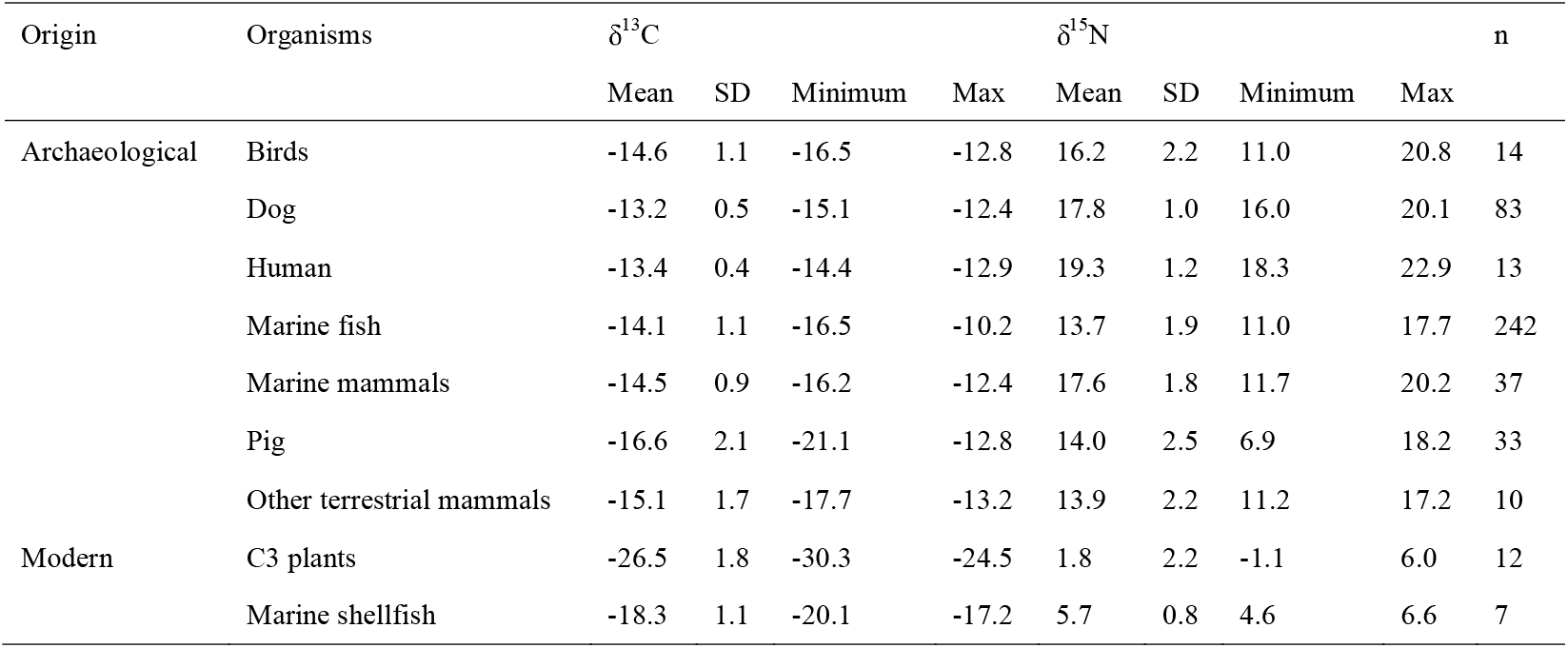
Summary of carbon and nitrogen stable isotope ratios of archaeological remains from the Hamanaka 2 site and modern food sources m the Rebun Island.

**Figure 1.**
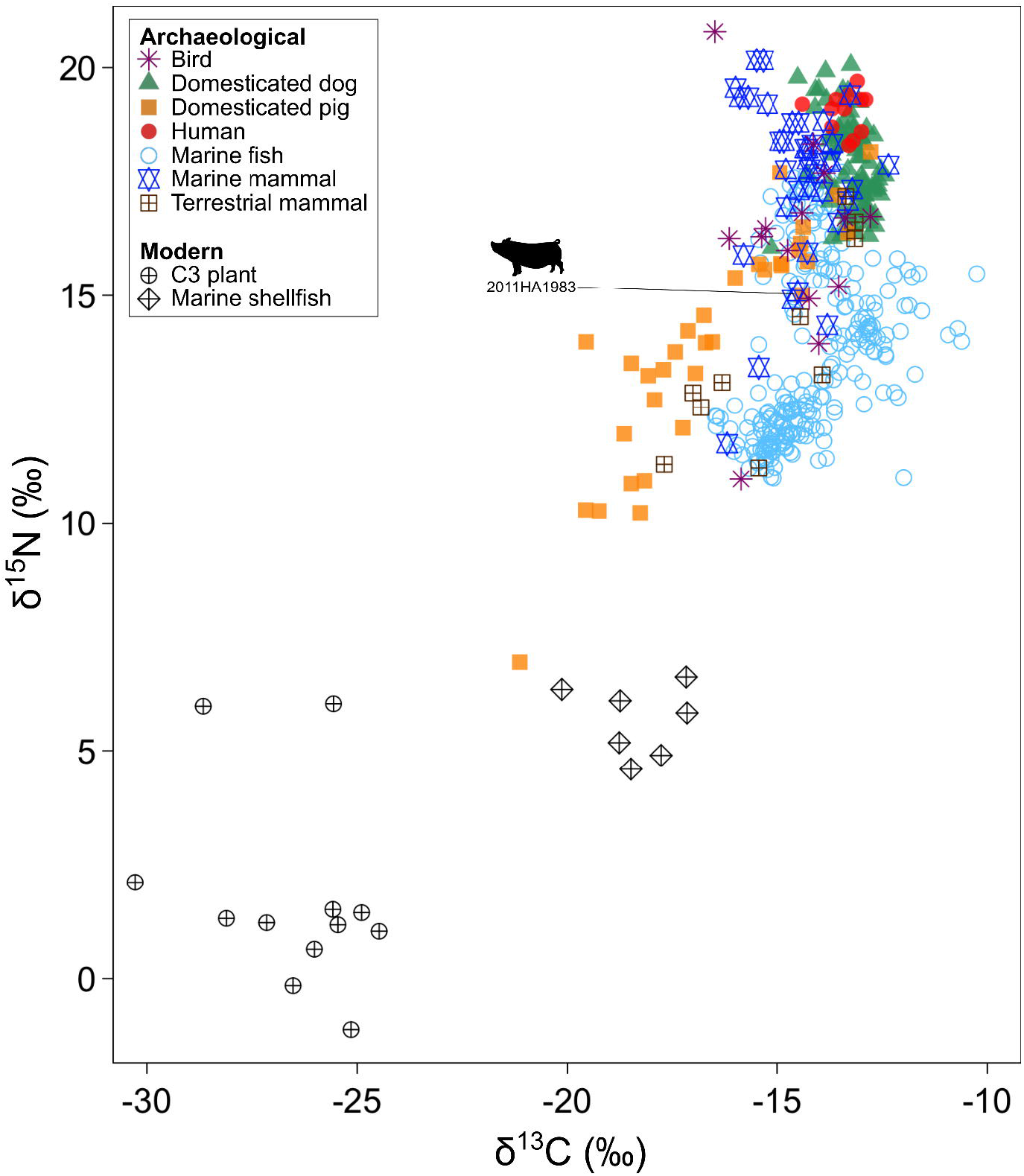
Carbon and nitrogen stable isotope ratios measured in archaeological skeletal collagen obtained from Hamanaka 2 and modern food sources of Rebun Island. The values of the pig (2022HA1983) on which the sequential analysis of enamel increments was performed are shown. Modern food sources are plotted with a +1.0‰ correction for the δ^13^C values considering the Suess effect (Friedli et al., 1986).

### Species differences in terrestrial consumers

The Bayesian mixing model package SIAR (Parnell et al., 2010) revealed that humans and dogs primarily relied on marine foods (≥90% of dietary protein), while pigs displayed variable mixed diets between terrestrial C3 plants and marine foods (Figure 2; Supplementary Table S6). Wild foxes (*Vulpes vulpes*) had an intermediate dietary protein contribution from terrestrial C3 plants (∼30%) (Figure 2; Supplementary Table S6), and their contribution was significantly greater in the Epi-Jomon period (Supplementary Figure S4; Supplementary Text S2). The dietary protein contributions from marine mammals were higher in humans (38.0–66.8% in the 95% credible interval [CI]) than in dogs (33.5–44.1% in 95% CI) (Figure 2; Supplementary Table S6). Human individuals have relatively homogenous isotope ratios among different sexes and age classes (Supplementary Table S7; Supplementary Figures S2 and S3). Dog individuals showed no significant diachronic trends (Supplementary Figure S4).

**Figure 2.**
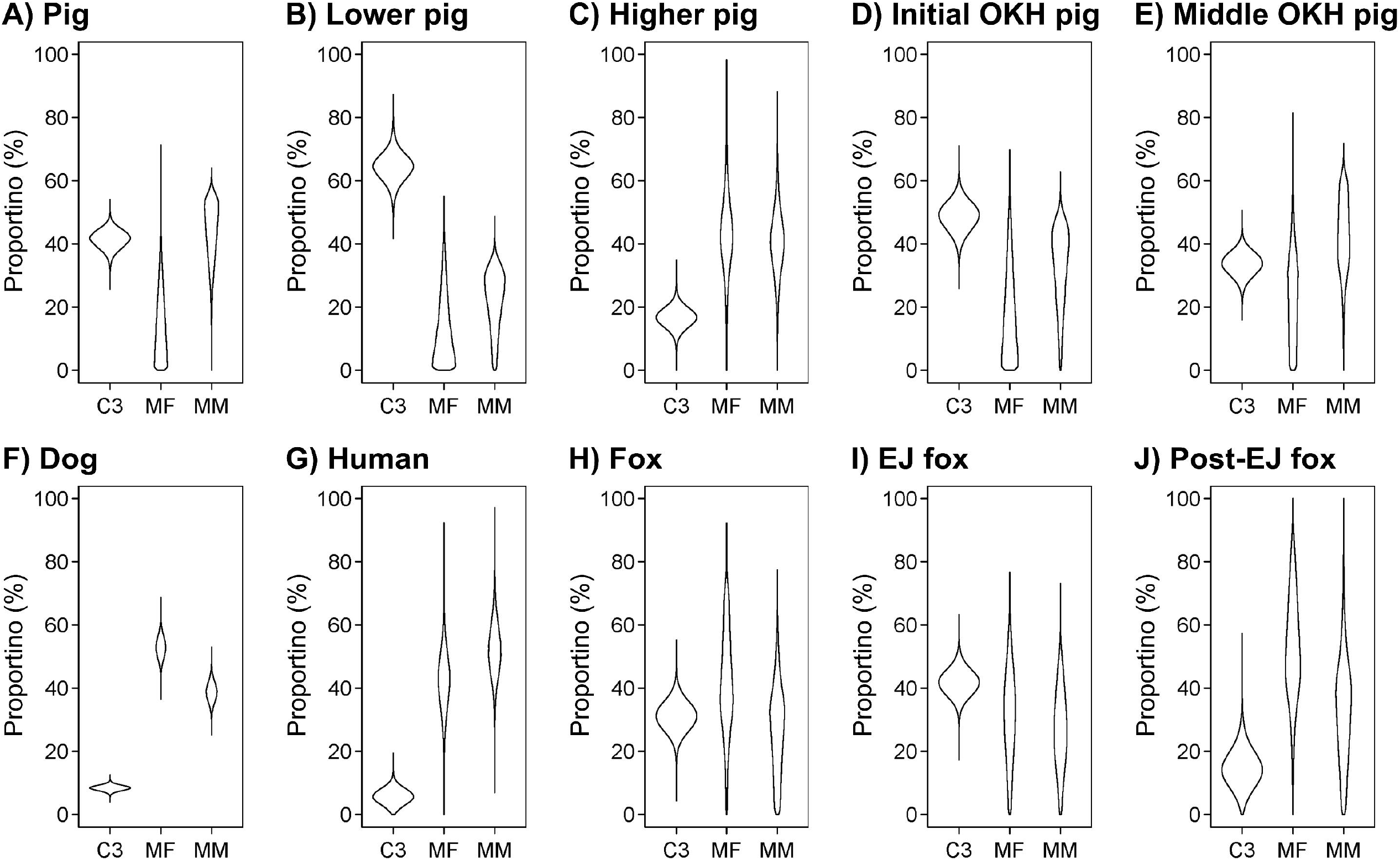
Beanplots showing the result of SIAR calculation for the proportions of protein contributions from each food source in different animals from Hamanaka 2. A) all pigs individuals; B) and C) each seven pig individuals with the lowest or highest δ^15^N values, respectively; D) and E) pigs from Initial or Middle Okhotsk periods, respectively; F), G), and H) all dogs, humans, and foxes, respectively; I) and J) foxes from Epi-Jomon or post Epi-Jomon (i.e., Okhotsk and Ainu) periods, respectively.

The stable isotope ratios of pig specimens ranged linearly between terrestrial and marine end members (from -21.1‰ to -12.8‰ in δ^13^C and 6.9‰ to 18.2‰ in δ^15^N) (Figure 1). The dietary protein contribution from C3 plants in pigs varied from 16.5% (11.5–21.7% in 95% CI) in seven individuals with the lowest δ^15^N values to 64.5% (56.6–71.9% in 95% CI) in those with the highest δ^15^N values (Figure 2; Supplementary Table S6). The dietary protein contribution from C3 plants in pigs decreased from 49.5% (40.7–56.1% in 95% CI) during the Initial Okhotsk period to 33.5% (27.3–39.4% in 95% CI) in the Middle Okhotsk period (Figure 2; Supplementary Table S6). While Mann–Whitney U-tests indicated no significant difference in pig δ^13^C values between these periods (1.4‰ lower in average, U = 69, p = 0.071), δ^15^N values were significantly lower in the Initial Okhotsk period (2.5‰ lower in average, U = 51, p = 0.011) (Supplementary Figure S4).

### Seasonal change in pig diet

To further explore the isotopic variation in pigs, sequential oxygen and carbon stable isotope analysis of tooth enamel was performed along the growth axis of the first and second incisors of a single pig specimen (2011HA1983). This pig individual showed intermediate to high δ^15^N values among the isotopic distribution of the Hamanaka 2 pigs (Figure 1). Since tooth enamel records chronological dietary changes during tooth growth and becomes isotopically inactive after maturation, this analysis can reveal dietary shifts throughout an individual’s life (Balasse, 2002; Balasse et al., 2006).

The oxygen (δ^18^O values) and carbon isotope ratios of enamel increments ranged from -9.0‰ to -4.5‰ and -11.4‰ to -7.0‰, respectively (Supplementary Table S8), displaying a clear inverse pattern (Figure 3). Using a bioapatite-diet spacing of +13.3 ± 0.3% (Passey et al., 2005), these δ^13^C values correspond to a diet exclusively composed of C3 plants (-24.7‰) and that of pigs with one of the lowest δ^13^C values of bone collagen (-20.3‰) (Figure 1). The pig’s diet exhibited lower δ^13^C values during the summer, corresponding to higher δ^18^O values and higher temperatures, and higher δ^13^C values during the winter, corresponding to lower δ^18^O values and lower temperatures (Figure 3). The maximum difference in incremental enamel δ^13^C values within this individual (4.4‰) accounted for approximately 53% of the maximum difference in δ^13^C values of bulk bone collagen among different individuals (8.3‰). These results indicate a cyclic seasonal shift in the proportion of terrestrial C3 plants and marine foods in the pig’s diet.

**Figure 3.**
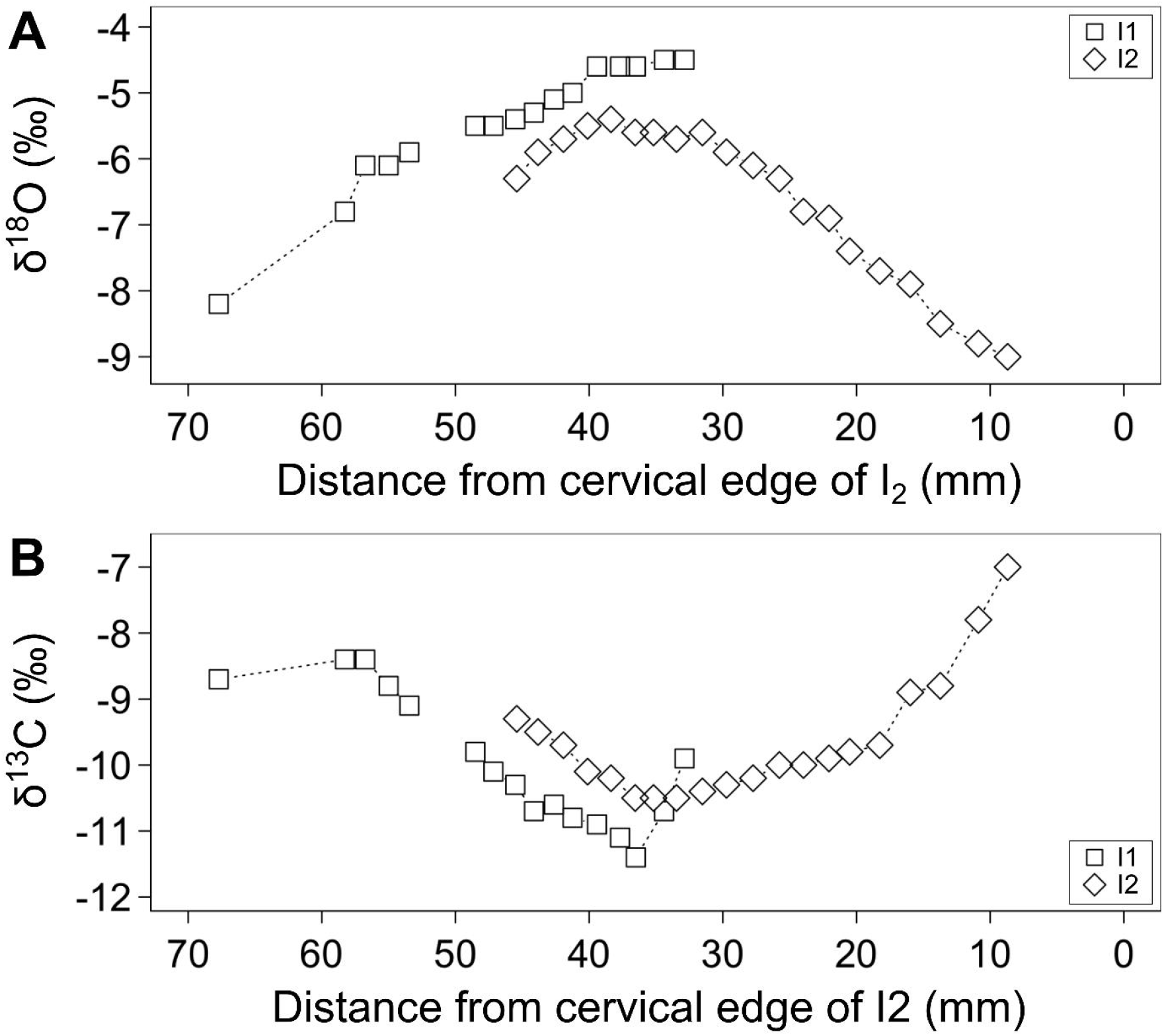
Oxygen and carbon stable isotope ratios measured in archaeological dental enamel increments sampled from pig 2022HA1983 of Hamanaka 2. The isotopic trajectories for I1 and I2 were aligned by referring to the previous results of Frémondeau et al. (2012).

## Discussion

### Feeding patterns and animal management

Stable isotope analysis revealed distinct feeding patterns and dietary differences among mammals, including humans, at Hamanaka 2 (Supplementary Text S2). Both humans and dogs showed a heavy reliance on marine foods, accounting for approximately 90% of their dietary protein intake. This result suggests dogs at Hamanaka 2 were fed by humans and appeared to be well cared for, as indicated by the lower frequency of early developmental stress markers compared to pigs (Onishi, 2015).

In contrast, pigs consumed a mix of terrestrial C3 plants and marine foods, with seasonal variation in their diet as revealed by sequential isotope analysis of tooth enamel (Figure 3). Given that bone collagen turnover rates are high during early life and that a significant proportion of pigs were slaughtered within a few years after birth (with individuals aged ≤2.5 years comprising 50–60%: Hattori, 2017), the bulk stable isotope ratios of pig bone collagen mostly represent the diet from the months immediately preceding death. During summer, pigs primarily consumed C3 plants, while in winter, they were likely fed marine foods, as a result of human intervention. Since bone collagen and enamel apatite primarily reflect isotopic signatures of protein and whole dietary macronutrients, respectively (Jim et al., 2004), the discrepancy between the dietary δ^13^C values of 2011HA1983 estimated from collagen and enamel was due to differences in carbon routing. This seasonal dietary shift suggests that pig management was influenced by the harsh subarctic climate of Rebun Island, where heavy snowfall in winter likely necessitated more intensive human husbandry. This aligns with zooarchaeological evidence suggesting that pigs were initially brought to the island and reared for a short period before slaughter during the Initial Okhotsk period, with a breeding population established by the Middle Okhotsk period (Hattori, 2017) (Supplementary Text S1). The stable isotope data corroborate this interpretation, showing a shift from a greater reliance on terrestrial foods, typical of Eurasian ancient pigs (Cucchi et al., 2016; Kuzmin et al., 2018), in the Initial Okhotsk period to an increased consumption of marine foods in the Middle Okhotsk period. However, pigs experienced greater stress, as indicated by the higher frequency of early developmental stress markers (Hattori, 2017).

### Failed invasion of pigs

Despite being a highly invasive species (Barrios-Garcia and Ballari, 2012; Risch et al., 2021), domesticated pigs introduced to Rebun Island during the Okhotsk period failed to establish a wild population, eventually disappearing from the archaeological record and the island. Analyzing this failed invasion through the frameworks of invasion ecology (Blackburn et al., 2011; Zenni and Nuñez, 2013), we identified three key factors that contributed to this failure: reduced propagule pressure, abiotic resistance due to the cold climate, and decreased resources (Figure 4).

**Figure 4.**
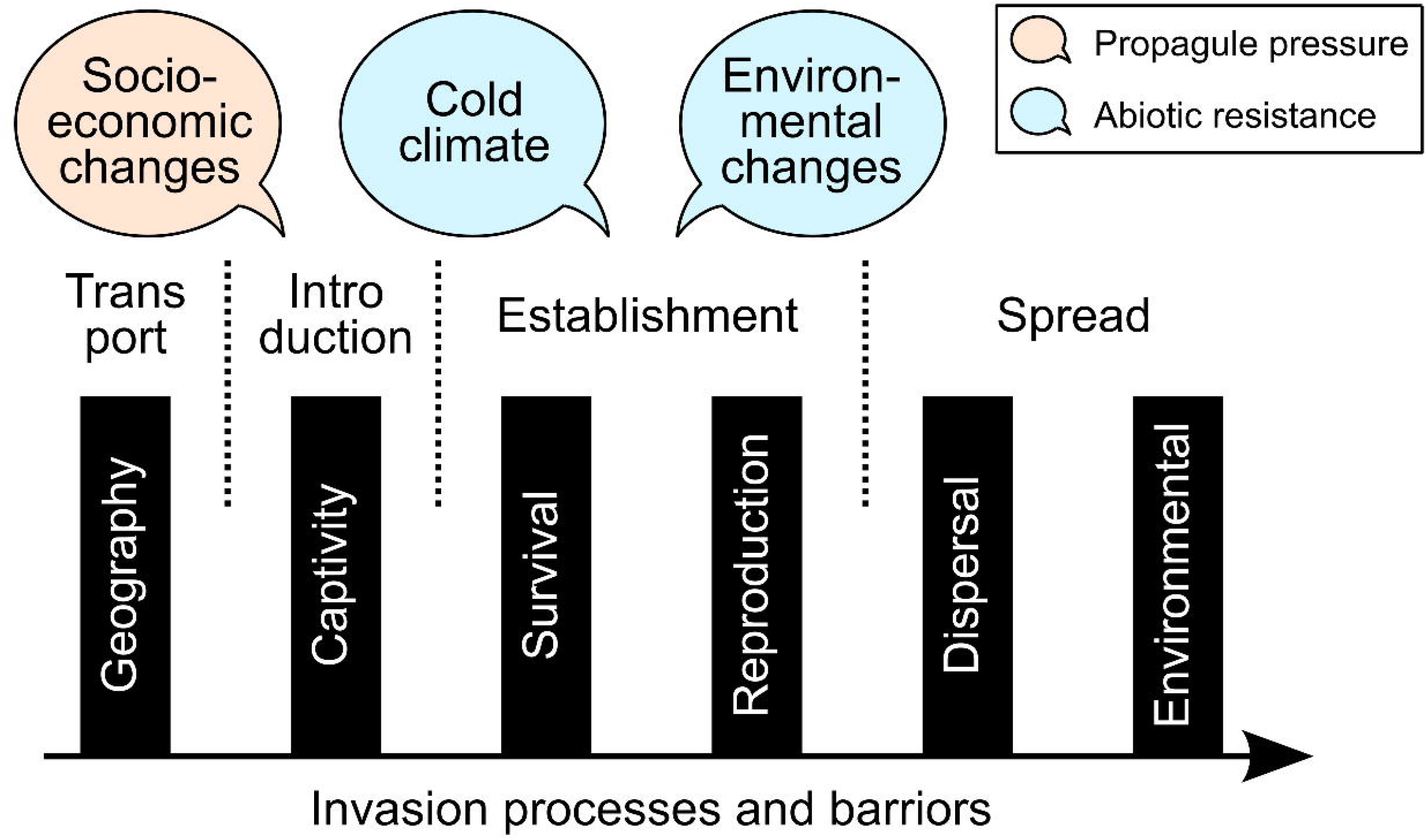
Schematic illustration of the frameworks of invasion ecology, with the three main factors that contributed to the failed invasion of archaeological pigs on Rebun Island.

First, socioeconomic changes in neighboring cultures surrounding the Okhotsk culture led to a decrease in the supply of pigs from northeastern Eurasia (Supplementary Text S3), which was the primary source for the Okhotsk people on Rebun Island (Watanobe et al., 2001). During the Initial and early Middle Okhotsk periods, the Okhotsk people traded extensively with the Mohe culture, which valued pigs as symbols of wealth (Minoshima, 2018). However, with the rise of the Bohai state and the subsequent decline in Mohe autonomy, trade routes were disrupted, reducing the availability of pigs (Minoshima, 2018, 2019; Usuki, 2005) and undermining the introduction stage of the invasion process (Figure 4). This decline in trade, coupled with the shift towards interactions with cultures in mainland Japan (Minoshima, 2018, 2019; Usuki, 2005), where pig husbandry was not prominent and pig meat consumption was prohibited due to Buddhism (Price and Hongo, 2020), likely reduced the motivation and ability to maintain pig populations on the island.

Second, the cold climate and heavy snowfall on Rebun Island posed significant challenges to pig survival and reproduction, reinforcing the barriers to establishment (Figure 4) (Supplementary Text S3). Pigs, which originated in Southeast Asia (Larson et al., 2005), are not naturally adapted to the subarctic conditions of Rebun Island (but see Lin et al., 2017), where winter temperatures are lower than their natural range (Oliver, 1993). Lower temperatures during winter are shown to be the strongest factor restricting the distribution of wild boars and pigs (McClure et al., 2015; Melis et al., 2006). The climate regime of the Medieval Little Ice Age (ca. 1550 cal AD) likely exacerbated these harsh conditions (Leipe et al., 2018). The sequential isotope analysis suggests that humans supplemented pigs’ diets with marine foods during winter, a diet that possibly caused nutritional stress due to its deviation from the pigs’ natural diet, which consists mostly of plants (Ballari and Barrios-Garcia, 2013). Additionally, pigs’ potential lack of adaptation to the high salt or arsenic contents of marine foods (Feldmann et al., 2000; Hall et al., 1975) and the physiological challenges posed by cold and snow (Danilov and Panchenko, 2012; Honda, 2009) further contributed to their inability to establish a stable population.

Third, climate change and increased human activity on Rebun Island during the Middle Okhotsk period further limited the availability of resources necessary for pig survival (Figure 4) (Supplementary Text S3). Pollen analysis from a sediment core at Lake Kushu, located 1–2 km east of Hamanaka 2, indicates a cooling trend throughout the Okhotsk period and a significant decrease in forest cover during the Middle Okhotsk period, likely due to increased clearance of woodland by humans (Leipe et al., 2018). This reduction in forest cover would have decreased the availability of terrestrial plant foods and suitable habitats for pigs (McClure et al., 2015; Melis et al., 2006), exacerbating the challenges they faced in surviving the harsh winter conditions, as well as the cooling climate. Although carred barley has been identified from Hamanaka 2 during the Okhotsk period (Leipe et al., 2017), archaeological evidence of horticulture in the Okhotsk culture is limited, and these crops could not be a major food source for pigs.

### Implications for contemporaneous invasion ecology

By investigating archaeological records, this study revealed that cold climates with heavy snowfall can substantially hinder the survival and reproduction of pigs, even with human intervention. While the negative impacts of pig invasions are particularly pronounced on islands (Risch et al., 2021), the case of Rebun Island during the Okhotsk period suggests that the limited terrestrial resources on small islands contributed to the failure of the pig invasion. Previous studies have demonstrated the difficulty of maintaining domesticated pigs on the smaller islands of prehistoric Polynesia and Palau, primarily due to the limited availability and diversity of resources (Clark et al., 2013; Giovas, 2006; Kirch, 2000). This implies that the success of pig invasions may vary depending on the climate and environmental conditions of the region. As climate change leads to warming, regions that were previously resistant to pigs because of the lower productivity may become more vulnerable to invasion (Melis et al., 2006).

This study represents a novel approach to studying failed invasions using archaeological data, offering a unique perspective on the long-term processes that influence invasion success or failure. By integrating archaeological records with contemporary invasion ecology frameworks, we can extend the time scales of invasion biology. Such an approach can also address knowledge gaps in the study of failed invasions, particularly for medium-to large-sized terrestrial animals, where contemporary examples are relatively rare (Zenni and Nuñez, 2013) but where skeletal remains are abundant in the archaeological record.

## Ethics

This work did not require ethical approval from a human subject or animal welfare committee. Research use of the archaeological remains was granted by the Free, Prior, and Informed Consent (FPIC) framework by the Ainu Association of Hokkaido.

## Data accessibility

Datasets related to this article are available in the Supplementary Information. Stable isotopic data (Tsutaya, 2026) have been uploaded to the IsoArcH repository (Salesse et al., 2018; Plomp et al., 2022).

## Declaration of AI use

AI tools were used to refine the initial version of texts in the Introduction and Discussion, but not to generate the contents. The final version of the texts was thoroughly edited by the authors.

## Author contributions

- Takumi Tsutaya: Conceptualization, Methodology, Formal analysis, Investigation, Writing (original draft), Writing (review & editing), Visualization
- Taichi Hattori: Methodology, Investigation
- Rin Onishi: Methodology, Investigation
- Chelsea E. Budd: Methodology, Investigation
- Hideki Minoshima: Methodology, Investigation, Writing (review & editing)
- Tomonari Takahashi: Methodology, Investigation
- Yu Hirasawa: Resources, Writing (review & editing)
- Sayaka Chiku: Formal analysis
- Takayuki Omori: Investigation
- Kohei Yamazaki: Investigation
- Minoru Yoneda: Resources
- Daisuke Kubo: Methodology, Investigation, Writing (review & editing)
- Hajime Ishida: Methodology, Investigation, Writing (review & editing)
- Takao Sato: Conceptualization, Methodology, Investigation, Resources
- Rick J Schulting: Conceptualization, Resources, Writing (review & editing)
- Hirofumi Kato: Conceptualization, Resources
- Andrzej W Weber: Conceptualization, Resources

## Acknowledgments

We thank the participants of the Rebun Archaeological Field School at Hamanaka 2.

