## Supplementary Information for "Leveraging archaeological specimens to investigate failed invasions: A case study on Okhotsk pigs, northern Japan"

**Supplementary Text S1. Archaeological context of the Hamanaka 2 site**

Rebun Island in Hokkaido is a relatively small island with an area of 81 $km^2$, and it has a history of continuous human occupation that spans from the Holocene Jomon hunter-gatherers to the historical Ainu populations (Table 1; Supplementary Figure S1). In this occupational sequence, pigs were introduced and domesticated only during the Okhotsk culture period (6th–13th centuries) and disappeared from the archaeological record thereafter (Table 1). The Hamanaka 2 site, the subject of this study, is a unique archaeological site where human occupation is continuously recorded and dated from the Jomon to the Ainu periods (Junno et al., 2021). During the Jomon and Epi-Jomon periods (299–258 cal BC and earlier), hunter-gatherer-fishers who used pottery similar to that of the Honshu mainland of Japan and displayed similar skeletal morphology, seasonally utilized Hamanaka 2 (Kato, 2015; Nomura and Utagawa, 2003). During the initial Okhotsk period (489–573 cal AD), the Okhotsk people, who migrated south from the Amur River basin through Sakhalin, appear in the records of the Hamanaka 2 site. The Okhotsk people had a distinct pottery type and skeletal morphology compared to the Jomon and Epi-Jomon peoples (Nomura and Utagawa, 2003). Ancient genome studies have also shown that they originated from the northern regions of the Eurasian continent (Sato et al., 2021). The Okhotsk people were hunter-gatherer-fishers who had more stable dwellings and heavily depended on marine resources (Naito et al., 2010; Tsutaya et al., 2014). The Middle Okhotsk period (573–850 cal AD) marked the peak of human activity, characterized by thick fish bone layers (Kato, 2015). Pollen analysis also indicates a significant reduction in forest cover on Rebun Island during this subperiod (Leipe et al., 2018). The Final Okhotsk period (850–1200 AD) marks a decline in human activity, during which the Okhotsk culture and people gradually merged with the Satsumon culture, the successor of Epi-Jomon culture, giving rise to the Ainu culture (Adachi et al., 2018; Nomura and Utagawa, 2003; Sato, 2019). At the Hamanaka 2 site, the historical Ainu period (1550–1900 AD) is characterized by continued subsistence practices of hunting, gathering, and fishing, but no evidence of active plant cultivation has been detected to date, similar to the previous periods (Kato, 2015; Leipe et al., 2017). Almost all the domesticated pig bones excavated from the site date back to the Okhotsk culture period, making a stark contrast with the abundant domesticated dog bones identified throughout the entire cultural sequence (Kato, 2015). This tendency of the failed invasion of pigs, in contrast to that of dogs, is the same when considering the entire Rebun Island (Hattori, 2017).

Previous zooarchaeological studies that examined the morphological characteristics of pigs and dogs excavated from Hamanaka 2 have revealed distinct patterns and temporal changes in animal management (Hattori, 2017; Onishi, 2015). The Number of Identified Specimens (NISP) of pigs shows that there were 5 specimens in the Initial Okhotsk period, 179 specimens in the Middle period, and 56 specimens in the Late period, indicating that pig populations were most abundant during the Middle Okhotsk period (Hattori, 2017). From the

Initial to the Middle Okhotsk period, the pigs became smaller and more stable in size, suggesting that a breeding population was established and stabilized, as well as the effect of insular dwarfism (Hattori, 2017). Additionally, the frequency of enamel hypoplasia (EH), which serves as a stress marker during early development, in pigs was higher at Hamanaka 2 (65.4%) compared to domesticated pigs from a contemporaneous Jomon site (Ubayama shell mound, Chiba) in Honshu (30.1%), with an increase in the occurrence of enamel hypoplasia from the latter part of the Middle to the Late Okhotsk period. The age structure of pigs during the latter Middle to Late periods was skewed towards juveniles, with the proportion of young pigs with unerupted second molars increasing from the early (~16%) to late (~44%) Middle Okhotsk periods. Previous ancient mitochondrial DNA analysis of domesticated pig skeletons from Kafukai A site in the Okhotsk period of the Rebun Island suggested recursive introduction of pigs from Sakhalin Island and the Amur River Basin of northeastern Eurasia (Watanobe et al., 2001). In agreement with this, the morphology of pig skeletons excavated from the Hamanaka 2 site resembled that of fully domesticated pigs in northeastern Eurasia, which were distinct from wild boars (Hattori, 2017).

Domesticated dogs raised on Rebun Island were more similar in size to dogs from northeastern regions of the Eurasian continent, rather than to Jomon dogs from Honshu, suggesting that the dogs at the Hamanaka 2 site were introduced from northeastern Eurasia via Sakhalin (Onishi, 2015). There were no significant morphological changes in dogs from the Hamanaka 2 site between the Epi-Jomon and Okhotsk periods (Onishi, 2015). Among the 125 dog individuals with an estimated age from the Hamanaka 2 site, there is a notably high proportion of infants [up to 4 months old] (53%) and subadults [4–8 months old] (26%) (Onishi, 2015). Enamel hypoplasia was observed in only one dog individual among hundreds of excavated dog remains (Onishi, 2015). During the Okhotsk period, spiral fractures of the femur were found in 17% of adult dogs (n = 84) and 11% of infants and subadults (n = 46), and cut marks on the humerus were identified in 4% of adult dogs (n = 85) and 2% of infants and subadults (n = 55), suggesting that dogs were consumed as a food source (Onishi, 2015).

**Supplementary Text S2. Materials and Methods**

Sample collection

Archaeological skeletal materials were obtained through the excavation campaigns of the Hamanaka 2 site from 2011 to 2016 (Kato and Iwanami, 2014; Kato and Naganuma, 2016, 2017; Kato et al., 2012, 2015). Seven human skeletons excavated from localities of the Hamanaka 2 site in previous excavations were also included in the analysis (Nishimoto, 2000; Hanihara et al., 1994; Ishida et al., 1994, 2002), as well as 6 individuals from the Nakatani location. Stable isotope ratios of bone collagen were newly obtained from a total of 33 pigs, 83 dogs, 11 humans, and 61 other faunal specimens. Previously reported carbon and nitrogen stable isotopic data on 2 human (Okamoto et al., 2016; Uchida-Fukuhara et al., 2024) and 242 fish (Tsutaya et al., 2018, 2022) bone samples from the Hamanaka 2 site were merged into the dataset of this study. Previously reported stable isotopic data on two human skeletons (Naito et al., 2010) were not included due to the lack of radiocarbon ages.

Bones excavated from layers VIII–VII, VI–IV, III, and II were assigned to the Epi-Jomon period and to the Initial, Middle, and Final phases of the Okhotsk period, based on the pottery typology and radiocarbon dating (Table 1; Junno et al. 2021). Specimens identified in situ were prioritized over specimens found from the dry sieving of the excavated soils. Fish bones were collected via water flotation using 9.52, 4, and 2 mm meshes (Tsutaya et al., 2022). Taxonomic identification was done by experienced physical anthropologists and zooarchaeologists, and samples for stable isotope analysis were selected from the identified bones.

Modern reference samples of plants and marine shellfish were collected from Rebun Island between summer and autumn 2016. Fruits of *Empetrum nigrum*, *Rosa rugosa*, *Vitis coignetiae*, and *Actinidia arguta*, whose seeds were excavated from the Hamanaka 2 site (Müller et al., 2016), were collected at the Botanical Garden for High-Altitude Plants or the Campsite of Midorigaoka Park under permission from the Hokkaido Regional Forest Office of the Forestry Agency, Japan. According to the caretaker, as far as they were aware, these plants had never been fertilized in the past until the time of sampling. Samples of the surf clam (*Pseudocardium sachalinense*), green sea urchin (*Strongylocentrotus droebachiensis*), and northern sea urchin (*Mesocentrotus nudus*) were purchased from local stores and sampled near Funadomari Bay, where the Hamanaka 2 site is located on its sand dune. When comparing the modern data with archaeological ones, a +1.0‰ correction for the $\delta^{13}C$ values was applied to correct the Suess effect (Friedli et al., 1986).

Carbon and nitrogen stable isotope analysis of bone collagen and modern samples

Carbon and nitrogen isotope analysis was performed in two different research institutions: the University of Tokyo and the University of Oxford. Samples analyzed in the former include all human remains and modern food samples, and those start with “ha2”, “hd”, or “hp” for their laboratory identifier, and the latter with “J” (Supplementary Tables S1–S5). The methodological details are described below. Stable isotopic data (Tsutaya, 2026) have been uploaded to the IsoArcH repository (Salesse et al., 2018; Plomp et al., 2022).

*Stable isotope analysis at the University of Tokyo*

Collagen was extracted from bone samples weighing 6–360 mg using the method described in Tsutaya et al. (2017, 2018). Briefly, bones were washed with water and then 0.2 M NaOH to remove exogenous matter and humic substances. Bone samples were then decalcified in 0.25 N or 0.5 N HCl at 4°C for 2–4 days. Some samples were washed with 0.2 M NaOH at 4°C for a few hours after decalcification to ensure the removal of humic substances. The decalcified samples were gelatinized and filtered using a glass fiber filter (Whatman GF/F) to remove noncollagenous substances. The filtered samples were then freeze-dried. Then, 350–500 μg of the resultant collagen was used for bulk carbon and nitrogen stable isotope analysis. Lipid extraction was not performed, and NaOH treatment was mostly performed before decalcification in this study, since a previous study demonstrated experimentally that these treatments do not affect the stable isotope ratios of collagen extracted from fish bones specifically at Hamanaka 2 (Tsutaya et al. 2018).

Edible parts were used for stable isotope analysis for modern food samples. Modern shellfish samples were freeze-dried and ground into a fine powder using a motor and pestle. Lipid extraction was performed on the crushed samples by soaking them in a chloroform:methanol (2:1 in volume) mixture and ultrasonicating for 10 mins. Lipid-extraction was repeated until the solvent became transparent. The lipid-extracted samples were washed with acetone and dried overnight. Modern plant samples were freeze-dried and ground into fine powder using a motor and pestle.

Carbon and nitrogen stable isotope ratios were measured using an elemental analyzer–isotope ratio mass spectrometry (Thermo Flash 2000 elemental analyzer, Finnigan ConFlo III interface, and Thermo Delta V mass spectrometer) at the University Museum, University of Tokyo, Japan. Approximately 0.4 mg of the samples was weighed into a tin capsule for the measurements. To measure the $\delta^{15}N$ values, up to 8 mg, depending on the nitrogen content of the samples, was additionally analyzed for modern plant samples. The $\delta^{13}C$ and $\delta^{15}N$ values were calibrated against the laboratory working standard (L-alanine: $\delta^{13}C = -19.6 \pm 0.2$‰; $\delta^{15}N = 8.7 \pm 0.2$‰) provided by SI Science (Saitama, Japan), whose values were determined by the NBS 19 and the International Atomic Energy Agency (IAEA) Sucrose ANU (calibrated against Pee Dee Belemnite and IAEA N1 and IAEA N2 (calibrated against AIR) international standards, respectively. Based on repeated measurements of the calibration standards, precision was determined to be less than ±0.1‰ standard deviation (SD) for both $\delta^{13}C$ and $\delta^{15}N$. The accuracy or systematic error was not determined (Szpak et al. 2017).

*Stable isotope analysis at the University of Oxford*

Archaeological faunal samples were prepared for stable isotope analysis using standard in-house protocols at the Stable Isotope Laboratory of the School of Archaeology, University of Oxford. Bone collagen was extracted using a modified version of the Longin (1971) method. This method involves using acid and $H_2O$ (Milli-Q) washes, freeze-drying, and isotope measurement on an Elemental Analyser linked to a continuous flow Sercon 20/22 dual inlet mass spectrometer. Instrument precision is approximately ±0.2‰ for both $\delta^{13}C$ and $\delta^{15}N$, which is based on repeated measurements of known standards. Known-value standards are included to first drift-correct and then calibrate the readings taken from the unknown samples. The samples were measured in duplicate, and an average value was taken. Isotopic ratios are calculated with reference to in-house standards, which for this project comprised alanine (−27.11‰ and −1.56‰ for $\delta^{13}C$ and $\delta^{15}N$, respectively), and in-house cow (−24.30‰ and 7.86‰) and seal (−12.54‰ and 16.14‰) bone collagen standards. The $\delta^{13}C$ measurements are reported on the VPDB scale, and the $\delta^{15}N$ measurements are reported with reference to AIR.

Quality control parameters are used in stable isotope analysis to ensure the accuracy of the data produced. These include a collagen yield of 1% or above at 5 mg or more, and a C/N ratio of between 2.9 and 3.6 (DeNiro 1985). The %C and %N of the collagen samples are also calculated, as this provides a reliable indicator of sample preservation (van Klinken 1999). For bone collagen, the ideal ranges for %C and % N are approximately 40–48% and 12–17%, respectively. The purpose of these parameters is to ensure that sufficient organic material (e.g., mainly collagenous protein) was preserved, and that this material retains an acceptable in vivo signal (as assessed by the C/N and collagen yield).

Sequential carbon and oxygen stable isotope analysis of dental enamel

A sequential carbon and oxygen stable isotope analysis was conducted on the tooth enamel of pig specimen 2011HA1894, which showed intermediate to slightly elevated $\delta^{15}N$ values among the isotopic distribution of the Hamanaka 2 pigs. Although all four mandibular incisors were fully erupted in this individual, enamel hypoplasia was observed approximately 1 to 10 mm from the crown of the first incisors. There was minimal evidence of tooth wear. The crown heights were 41.92 mm for the first mandibular incisor ($I_1$) and 47.96 mm for the second mandibular incisor ($I_2$) analyzed. The dentin root of both $I_1$ and $I_2$ was not fully formed. The enamel sampling method followed the protocol described in a previous study (Frémondeau et al., 2012). First, the surface cementum was removed using a drill, exposing the enamel surface. Then, using a dental drill equipped with a tungsten carbide bur (Jet carbide burs, SHOFU Inc.), enamel powder was extracted at 1.5 mm intervals along the growth direction of the tooth by removing 1 mm-wide strips of enamel. Approximately 3 mg of enamel powder was collected from each sampling strip. Sampling was not performed in areas of the first incisor where enamel hypoplasia was identified at the tip. In total, 18 enamel powder samples were collected from the first incisor and 20 samples from the second incisor.

The enamel powder samples were soaked in 2.5% sodium hypochlorite for 24 hours to remove organic matter, followed by rinsing with Milli-Q water. Then, the samples were immersed in a 0.1 M acetic acid solution for 4 hours to remove exogenous carbonates, and then rinsed again with Milli-Q water. The chemically washed samples were dried at 70°C for 24 hours in an oven and then ground to a uniform grain size using an agate mortar and pestle.

Carbon and oxygen isotope ratios of enamel were measured by using a Kiel device coupled with an isotope ratio mass spectrometer (Kiel Device and MAT 253, Thermo Fisher Scientific) at the Department of Geology, National Museum of Nature and Science. The $\delta^{13}C$ and $\delta^{18}O$ values were calibrated against international standards NBS19 and NBS18. The measurement precision (standard deviation) for NBS19 was ±0.01‰ for carbon isotopes and ±0.04‰ for oxygen isotopes. The isotopic trajectories for $I_1$ and $I_2$ were combined in Figure 3, as previous results showed that the 27 mm point from the cervical edge of I1 corresponds to the same time point as the crown of $I_2$ (Frémondeau et al., 2012).

Radiocarbon dating

Approximately 2.5 mg of extracted collagen was combusted into carbon dioxide. After combustion, the purified carbon dioxide was introduced into a vacuum glass line and sealed, along with hydrogen gas (2.2 times the molar amount of carbon) and approximately 2 mg of pre-weighed iron catalyst, into a reaction tube. The reduction was carried out by heating the tube at 650°C for 6 hours, resulting in the formation of graphite. Radiocarbon ages were measured by using accelerator mass spectrometry at the University Museum, the University of Tokyo.

Radiocarbon ages were calibrated using the OxCal software (Bronk Ramsey, 1995) and atmospheric and marine data sets (IntCal20 and Marine20, respectively; Reimer et al., 2020; Heaton et al., 2020). Considering the marine-based diet of the Okhotsk people (Naito et al., 2010; Tsutaya et al., 2014), the contribution from marine carbon was estimated to be 94.5 ± 2.0%, based on the results of SIAR analysis. Three types of the local marine reservoir effects measured for Soya Current (⊿R = -64 ± 36 at Otaru, Hokkaido, recalculated with Marine20), for East Sakhalin Current (⊿R = 260 ± 35 at Svobodnaya, Sakhalin, recalculated with Marine20), and for in-between (⊿R = 217 ± 35 at Gulf of Aniva, Sakhalin, recalculated with Marine20) affecting Rebun Island were used for correction (Yoneda et al., 2007). Based on the correspondence between the age of the archaeological layers and the human remains from the Nakatani location, the calibrated radiocarbon ages were relatively younger when using the local marine reservoir effects for the Soya Current and older when using the local marine reservoir effects for the East Sakhalin Current, respectively (Supplementary Table S1). The calibrated radiocarbon age, taking into account the local marine reservoir effects, agreed better with the age of the archaeological layers (Supplementary Table S1). To achieve a more accurate calibration, information about the ancient fishing place and the contribution of different ocean currents to the place is needed.

Data analysis

The isotopic results were analyzed and visualized by using R software, version 4.2.3 (R Core Team, 2023).

Dietary protein contributions from different food categories were calculated using the R package siar, version 4.2 (Parnell et al., 2010). Dietary items were classified into terrestrial C3 plants (C3), marine fish (MF), and marine mammals (MM), considering archaeological evidence suggesting that wild terrestrial animals made a minimal dietary contribution to the consumer diet in Hokkaido prehistory (Naito et al., 2010; Tsutaya et al., 2014). Carbon stable isotope ratios of modern plant samples with the correction of +1.0‰ for the Suess effect (Friedli et al., 1986) were used. Isotopic offset from prey collagen to consumer collagen was set as 1.5 ± 0.5‰ for $\delta^{13}C$ values and 4.0 ± 1.0‰ for $\delta^{15}N$ values (Lee-Thorp, 2008). Isotopic offset from prey plants to consumer collagen was set as 5.0 ± 0.5‰ for $\delta^{13}C$ values and 4.0 ± 1.0‰ for $\delta^{15}N$ values (Lee-Thorp, 2008). Because dogs and marine fish showed almost identical $\delta^{13}C$ and $\delta^{15}N$ values, it is possible that the contribution from marine fish would be overestimated due to the possible consumption of domesticated dogs by humans. Additionally, terrestrial invertebrate animals, which are important dietary sources for pigs, typically exhibit higher $\delta^{13}C$ and $\delta^{15}N$ values than terrestrial C3 plants (Hyodo et al., 2011). However, due to the unavailability of such samples, we have not included terrestrial invertebrate animals in our analysis, and terrestrial C3 plants were used as a representative of terrestrial dietary sources. Therefore, the calculated terrestrial dietary contribution in pigs would be overestimated to a certain degree.

### Supplementary Text S3. Stable isotopic results of humans, dogs, and foxes

Humans had relatively homogenous $\delta^{13}C$ and $\delta^{15}N$ values with little systematic difference in sex, age class, and chronological period (Supplementary Figure S2 and S3). Mann-Whitney U-tests showed no significant difference between females and males in $\delta^{13}C$ ($U = 17.0$, $p = 0.574$) and $\delta^{15}N$ ($U = 14.5$, $p = 0.926$) values in humans. Mann-Whitney U-tests showed no significant difference between non-adults and adults in $\delta^{13}C$ ($U = 8.5$, $p = 0.067$) and $\delta^{15}N$ ($U = 33.0$, $p = 0.300$) values in humans. No significant correlation was found between the calibrated calendar age and isotope ratios (Spearman's rank sum correlation test, $\delta^{13}C$: $S = 226.1$, $p = 0.202$, $\delta^{15}N$: $S = 355.9$, p-value = 0.943).

Chronological dietary differences were observed in wild foxes but not in domesticated dogs at the Hamanaka 2 site (Supplementary Figure S4). The Kruskal–Wallis test did not reveal significant chronological differences in $\delta^{13}C$ ($X^2 = 6.65$, $p = 0.084$) and $\delta^{15}N$ ($X^2 = 3.89$, $p = 0.274$) values of dogs. However, Mann-Whitney U-tests revealed significantly higher $\delta^{13}C$ ($U = 0.0$, $p = 0.016$) and $\delta^{15}N$ ($U = 1.0$, $p = 0.032$) values in foxes during the Okhotsk and Ainu periods compared to the Epi-Jomon period.

Although both humans and dogs demonstrated a heavy reliance on marine foods, there was a clear dietary niche differentiation between them. Humans primarily consumed marine mammals, while dogs mainly fed on marine fish (Figure 2, Supplementary Table S6). This finding is consistent with previous studies of the Okhotsk populations at the Moyoro site in eastern Hokkaido (Tsutaya et al., 2014). Notably, the isotope ratios of dogs were nearly identical to those of marine mammals, suggesting the possibility that humans may have also consumed dogs, as supported by zooarchaeological evidence (Onishi, 2015). Future analyses of ancient biomolecules in human dental calculus could clarify this possibility (Uchida-Fukuhara et al., 2024).

Foxes, in contrast, showed a significant increase in stable isotope ratios during the Okhotsk and Ainu periods, characterized by increased human activities and the formation of thick fish bone layers in the former, compared to the Epi-Jomon period, indicating an increased consumption of marine foods (Figure 2; Supplementary Figure S4). Although foxes were not domesticated, they are commensal to humans, scavenging on waste from human food sources (Contesse et al., 2003). Therefore, the dietary habits of foxes can reflect the anthropogenic impacts on the surrounding environments in archaeological settings, as has been shown in Paleolithic sites in Europe (Baumann et al., 2020a, 2020b).

**Supplementary Text S4. Applying the frameworks of invasion ecology**

In invasion ecology, a unified framework is commonly used to divide the invasion process into four stages: transport, introduction, establishment, and spread (Blackburn et al., 2011). To progress to the next stage for an invasive population, various barriers such as survival and reproduction must be overcome (Blackburn et al., 2011). Additionally, five categories that influence the success or failure of invasions have been proposed: propagule pressure, abiotic resistance, biotic resistance, genetic constraints, and mutualist release (Zenni and Nuñez, 2013). Following these frameworks, we identified three key factors that led to the failure of the archaeological pig invasion on Rebun Island, based on the results of stable isotope analysis, as well as previous findings from zooarchaeology, archaeology, and history.

First, socioeconomic changes in neighboring cultures surrounding Okhotsk culture lowered the propagule pressure of pigs, which strengthened the captivity barrier and led to the failure of introduction after the Middle Okhotsk period. Pigs associated with the Okhotsk culture on Rebun Island originated from northeastern Eurasia (Watanobe et al., 2001). A breeding population was probably established on the island during the Middle Okhotsk period (Hattori, 2017). It is assumed that the supply of live pigs from Eurasia remained an important source of propagule pressure, especially when the local population experienced sudden, accidental declines. However, historical and archaeological evidence suggests that the availability of pigs from Eurasia significantly decreased or even ceased entirely in the latter half of the Middle Okhotsk period. During the Initial and early Middle Okhotsk periods (6th–7th centuries), the Okhotsk people engaged in active trade with the population known as the Mohe, who inhabited present-day Jilin and Heilongjiang provinces in China, as well as the Russian Far East (Amur Basin and Primorye) (Usuki, 2005). Pigs were obtained through these trade networks (Minoshima, 2018). Isotopic analysis and historical records indicate that the Mohe were primarily agriculturalists who cultivated C4 plants (Minagawa, 2002) and lived in a stratified society with officials and local leaders (Usuki, 2005), where pigs were considered symbols of wealth (Minoshima, 2018). However, in the latter half of the Middle Okhotsk period, around the mid-8th to early 9th centuries, the Mohe were dominated by the Bohai state, which was established at the end of the 7th century (Minoshima, 2018, 2019; Usuki, 2005). This conquest restricted the Mohe's ability to engage in free trade (Minoshima, 2018, 2019; Usuki, 2005). Consequently, the Okhotsk culture on Rebun Island reduced its trade with Eurasia and increased its interactions with the Satsumon culture on mainland Hokkaido and historical cultures on Honshu (Minoshima, 2018, 2019; Usuki, 2005). This shift in trading partners is clearly reflected in the changing array of exotic artifacts found at Okhotsk sites (Minoshima, 2019). After the fall of Bohai in AD 926, the Jurchen, the successor of Mohe, resumed autonomous trade, and Eurasian-origin artifacts began to reappear in Okhotsk cultural sites during the Final Okhotsk period in the 10th century and beyond (Minoshima, 2019). However, during the same period on the Honshu mainland of Japan, pig husbandry was minor, and the consumption of four-legged animals, including pigs, was strictly avoided due to Buddhist beliefs (Price and Hongo, 2020). Thus, while pigs were probably valued and obtained as symbols of wealth from the Mohe during the Initial Okhotsk period, the connection to this supply source ceased in the later Middle Okhotsk period. Simultaneously, trade intensified with Honshu cultures that did not value pigs or even avoided them. These socio-economic changes likely reduced both the availability of pigs and the motivation to obtain them, leading to a decline in propagule pressure among the Okhotsk pigs on Rebun Island.

Second, the physiological constraints of pigs faced abiotic resistance of the cold climate and snowfall, which strengthened survival and reproduction barriers and led to the failure of establishment in Rebun Island. Lower temperatures during winter are shown to be the strongest factor restricting the distribution of wild boars and pigs (McClure et al., 2015; Melis et al., 2006). Domesticated pigs originated from Southeast Asia (Larson et al., 2005), and Rebun Island is located even further north than the natural range of wild boars (Oliver, 1993), where the cold temperatures during winter are unfavorable for pigs. The Okhotsk period corresponds to the Medieval Little Ice Age regime, and the climate was likely even colder than it is today (Leipe et al., 2018). Snowfall is also an unfavorable climatic factor for pigs. Wild boars tend to avoid habitats in regions with deep snow (Honda, 2009), as it becomes difficult for them to move when the snow depth exceeds 40 centimeters (Lozan, 1995). Large-scale snowfall can sometimes lead to mass mortality (Danilov and Panchenko, 2012). Pigs are omnivorous but prefer plant foods over animal foods (Ballai and Barrios-Garcia, 2013). In the typical diet of wild boars, plant matter constitutes more than 90% of their intake (Ballai and Barrios-Garcia, 2013). During winter on Rebun Island, snowfall likely restricted access to terrestrial C3 plants. As suggested by sequential isotope analysis of enamel (Figure 3), humans may have supplemented the pigs' diet with marine resources. The practice of feeding fish to domesticated pigs by the Nanai people in the mid-19th century in the Ussuri River is recorded in an expedition note of the time (Maak, 1859). This diet, dominated by marine products, deviated significantly from pigs' natural diet and likely caused nutritional stress. In the Orkney Islands of Scotland, sheep that survive on seaweed during winter have been known since the Neolithic period (Balasse et al., 2006; Schulting et al., 2017). However, this diet requires various physiological adaptations, such as tolerance to high salt concentrations (Hall et al., 1975), resistance to high arsenic levels (Feldmann et al., 2000), and adaptation to low bioavailability of copper (MacLachlan and Johnston, 1982). Such adaptations would require hundreds of years, and it remains uncertain whether pigs could achieve similar adaptations. The cold climate, thick snow cover, and restricted terrestrial foods on Rebun Island in winter posed significant barriers to pig survival and reproduction. The high frequency of EH observed in pig teeth suggests that the pigs were under substantial nutritional stress even with human care (Hattori, 2017).

Third, climate change and increased human activity in Rebun Island posed further abiotic resistance to resource availability, which strengthened survival and reproduction barriers and led to the failure of establishment in Rebun Island during the Middle Okhotsk period. Previous pollen analysis from a sediment core at Lake Kushu, located 1–2 km east of Hamanaka 2, reveals that during the Middle Okhotsk period, corresponding to AD 370-850, forest cover significantly decreased (Leipe et al., 2018). As forests are important sources of plant foods (Melis et al., 2006) and woody materials for winter-time nests during snowfalls (Danilov and Panchenko, 2012) in pigs, this reduction would exacerbate the already poor food availability and winter-time living conditions, making their survival and reproduction even more difficult. The decrease in forest cover is likely attributed to increased human activity, particularly logging (Leipe et al., 2018). However, the simultaneous increase in both pig remains and the frequency of EH during the Middle Okhotsk period (Hattori, 2017) suggests a potential negative feedback: the pigs' increased foraging and rooting behaviors may have contributed to the reduction in forest cover, thereby reducing their own food sources and increasing their stress levels. In this scenario, although the pigs ultimately failed to successfully invade the island, they may have had a destructive impact on the local vegetation in the past. Furthermore, the previous results of a consistent decrease in temperate deciduous tree pollen during the Okhotsk period indicated a cooling trend on Rebun Island under the Little Ice Age regime (Leipe et al., 2018). Such climate change in the past is an abiotic factor that further suppressed the establishment of pigs, which originated in Southeast Asia.

Although they did not have a substantial effect on the failed invasion, other categories that influence the success or failure of invasions can also be evaluated in the case of Okhotsk pigs in Rebun Island. Regarding biotic resistance, it can be argued that Rebun Island was actually a favorable environment for pig invasion. The island had few medium- or large-sized terrestrial vertebrates, with native species like foxes and martens, and lacked strong predators or mammals occupying the same ecological niche as pigs (Garza et al., 2018). However, predation pressure from humans and domesticated or feral dogs introduced by humans might have harmed the pigs' survival. Concerning genetic constraints, the pigs introduced to Rebun Island might have been well-suited to the cold climate (Lin et al., 2017). These pigs were likely fully domesticated in the northeastern part of Eurasia (Hattori, 2017; Watanobe et al., 2001), suggesting they may have already undergone genetic or physiological adaptations to cold environments by the time they arrived on the island. To explore this hypothesis further, ancient genome analysis of pigs from Eurasia and Rebun Island would be necessary. As for mutualist release, the situation might have been more favorable to invasion. Humans, who provided care and feeding during the winter, were crucial mutualists for the pigs. However, at Hamanaka 2, there is evidence that human activity diminished during the transitional periods between the Epi-Jomon and Okhotsk periods, and between the Okhotsk and Ainu periods (Junno et al., 2021; Kato, 2015). If such trends were common across Rebun Island, the near absence of humans during these times may have negatively impacted the pigs' ability to establish themselves. A more detailed examination would require a quantitative reconstruction of changes in human activity across the entire island (Leipe et al., 2018; Müller et al., 2016).

**Supplementary Figures**

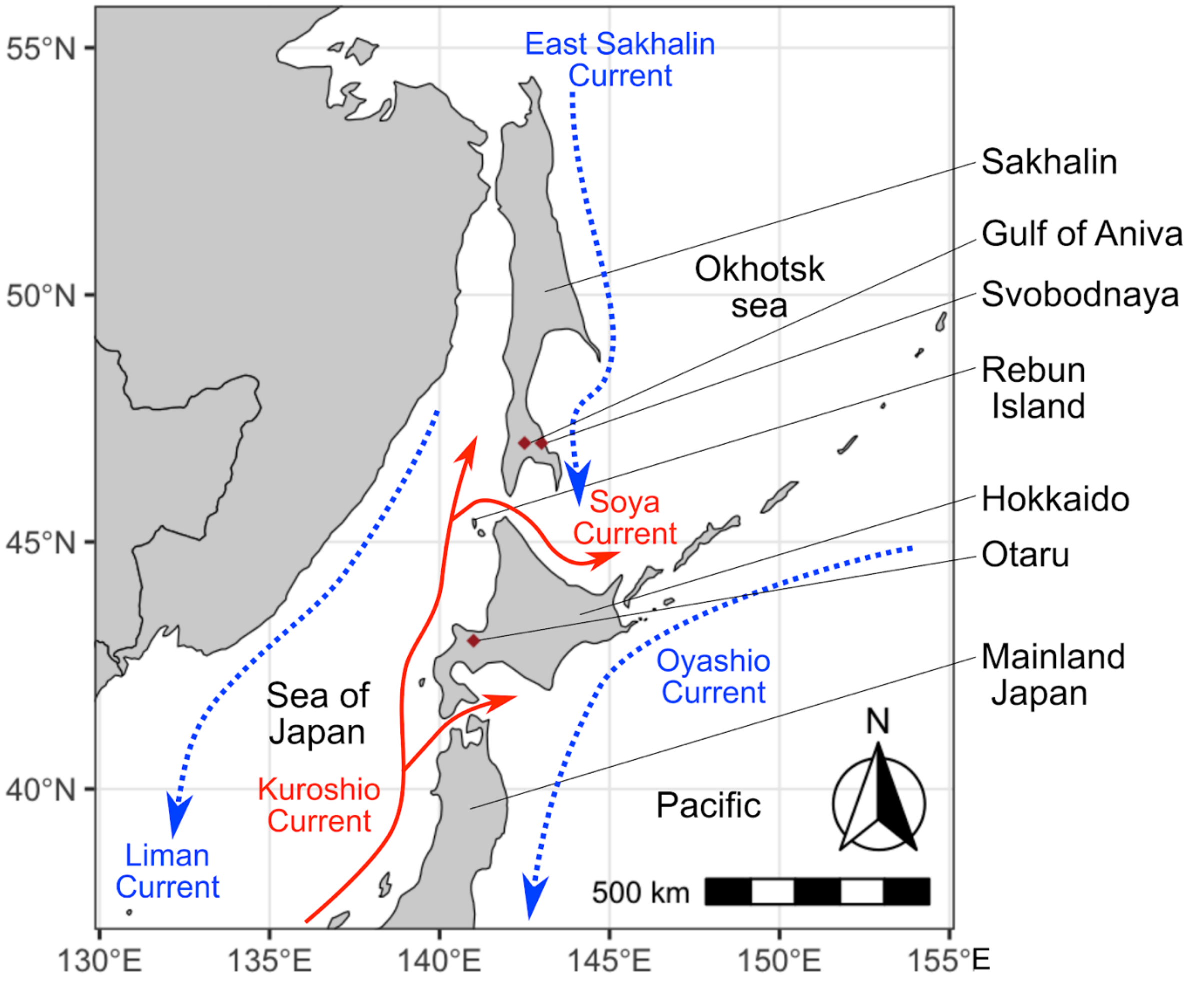

**Supplementary Figure S1.** Map showing the location of Rebun Island, major currents, and previously reported sites for the calculation of local $^{14}C$ marine reservoir effects.

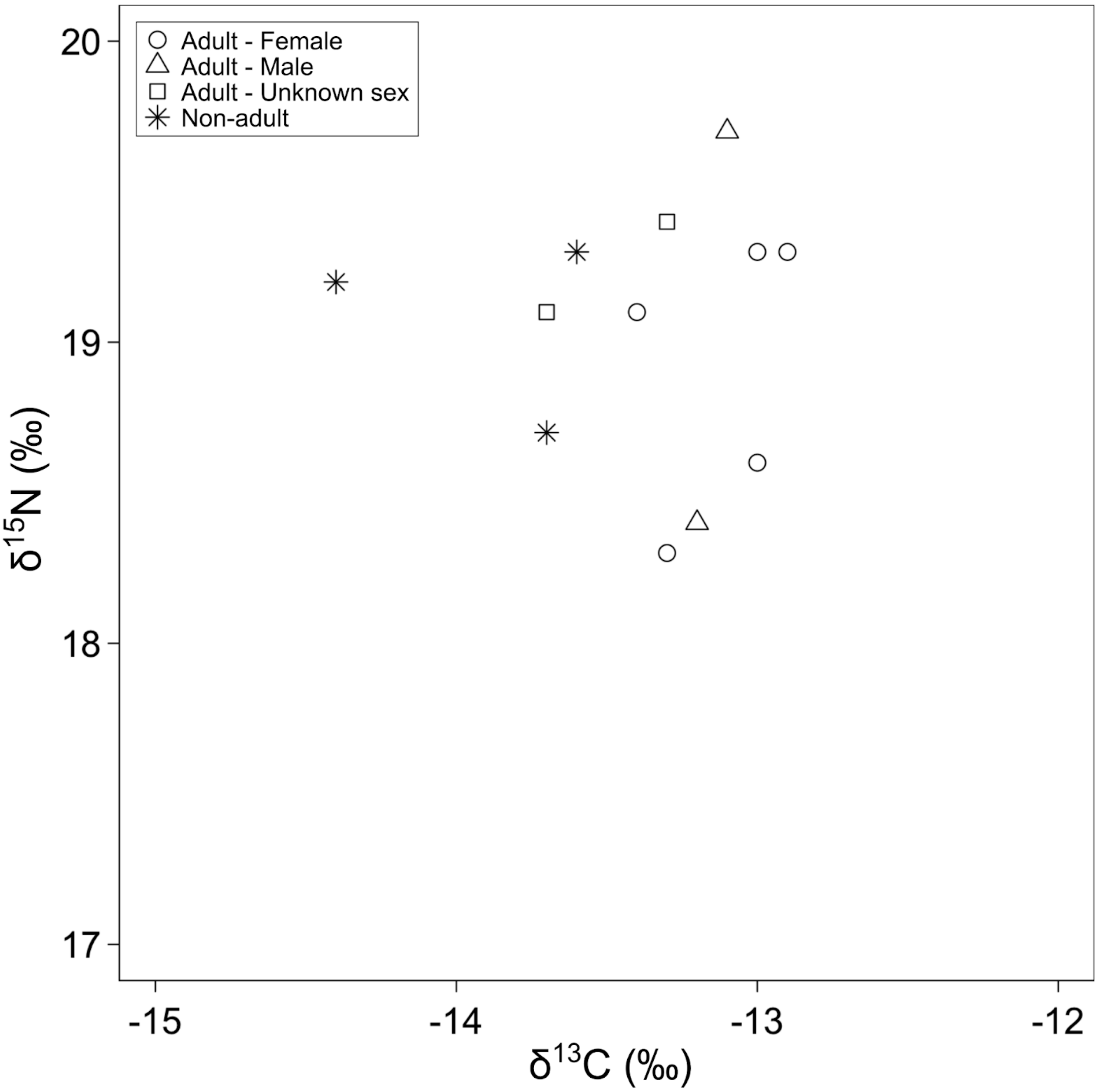

**Supplementary Figure S2.** Carbon and nitrogen stable isotope ratios of human skeletons from the Hamanaka 2 site.

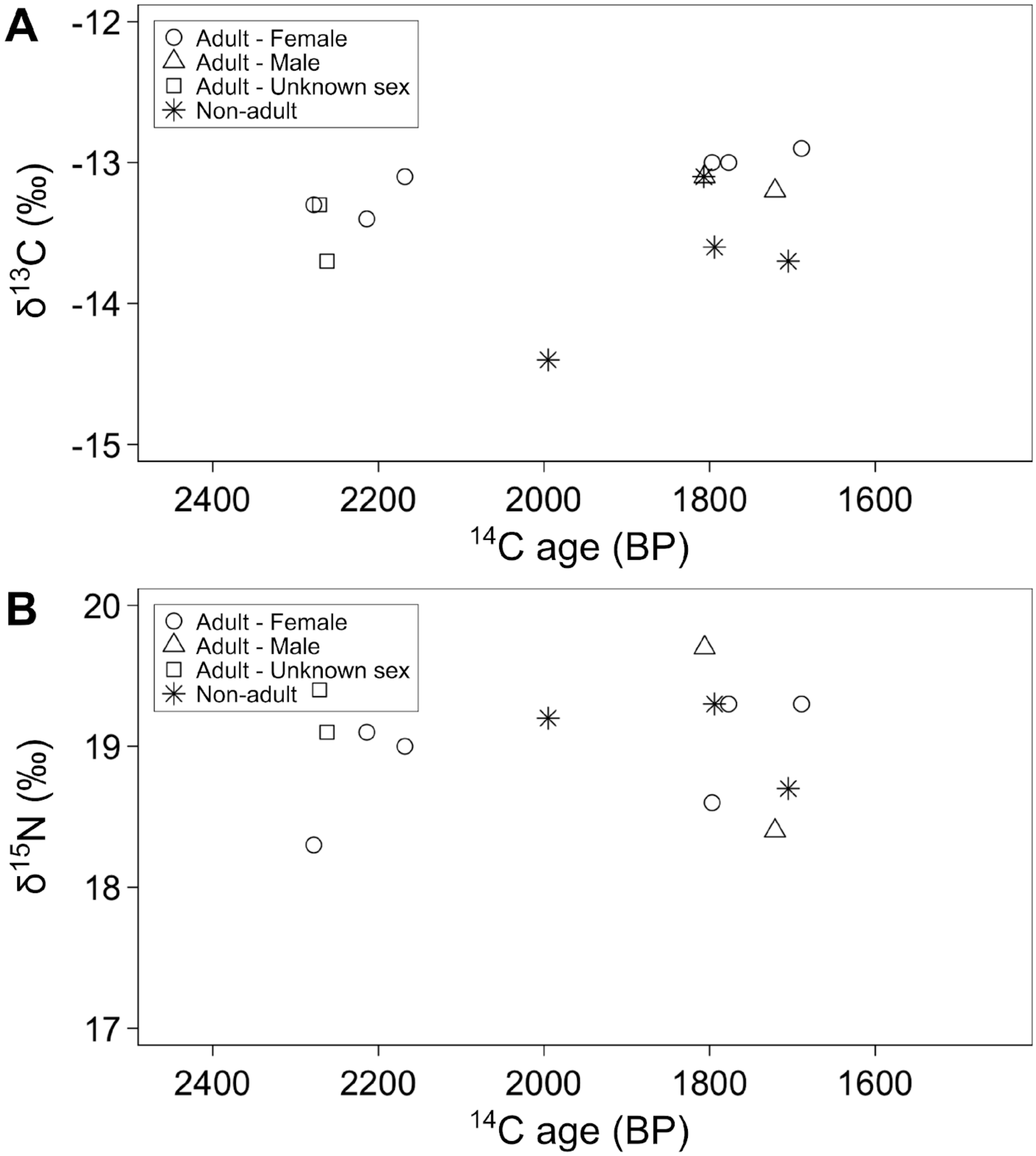

**Supplementary Figure S3.** Relationship between $^{14}C$ age and **A)** carbon or **B)** nitrogen stable isotope ratios of human skeletons from the Hamanaka 2 site.

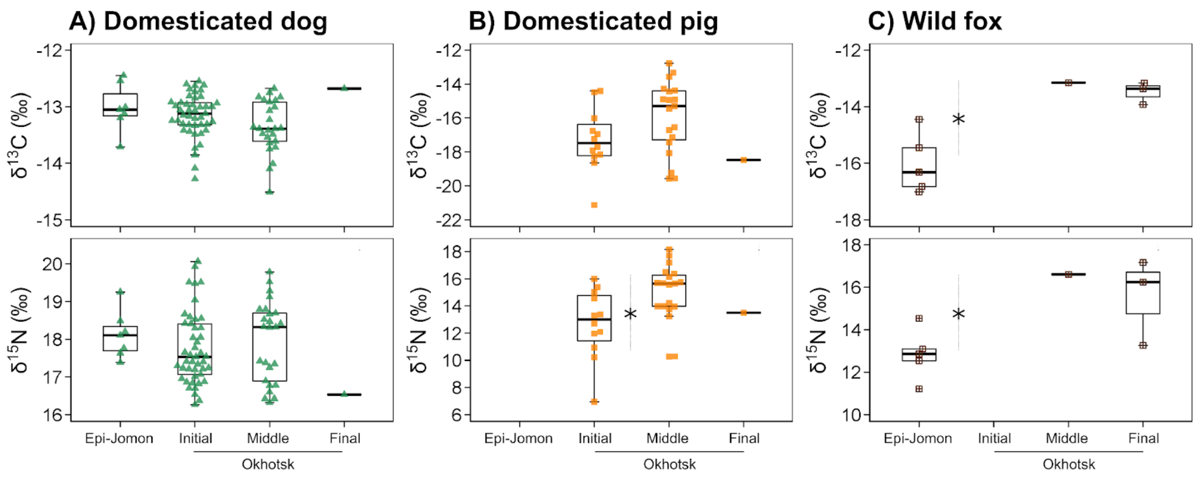

**Supplementary Figure S4.** Chronological difference of carbon and nitrogen isotope ratios of **A)** domesticated dogs, **B)** domesticated pigs, and **C)** wild foxes from the Hamanaka 2 site. Significant chronological differences are shown with a dotted bar with an asterisk.

### Supplementary Tables

**Supplementary Table S1.** Stable isotopic data of human skeletons from the Hamanaka 2 site. Calibrated $^{14}$C ages with different references for local marine reservoir effects are shown.

| ID | Element | %C | %N | $\delta^{13}$C | $\delta^{15}$N | C/N | Location | Layer | Period | Reference |
|---|---|---|---|---|---|---|---|---|---|---|
| 93-1 | Rib | 44.7 | 16.7 | -13.1 | 19.7 | 3.1 | – | – | Middle Okhotsk | This study |
| HA2 NAT002 | Rib | 41.8 | 15.0 | -12.9 | 19.3 | 3.2 | Nakatani | IIb | Final Okhotsk | Okamoto et al. (2016) |
| HA2 NAT003 | – | 45.4 | 16.3 | -13.6 | 19.3 | 3.2 | Nakatani | IIIb | Middle Okhotsk | This study |
| HA2 NAT001 | Rib | 47.0 | 16.6 | -14.4 | 19.2 | 3.3 | Nakatani | IIIe | Middle Okhotsk | This study |
| 2017HA0997 | Rib | 40.7 | 14.7 | -13.3 | 19.4 | 3.2 | Nakatani | V | Initial Okhotsk | This study |
| 2017HA0999 | Vertebra | 46.2 | 16.7 | -13.7 | 19.1 | 3.2 | Nakatani | V | Initial Okhotsk | This study |
| HA2 NAT004 | Rib | 40.8 | 15.0 | -13.3 | 18.3 | 3.2 | Nakatani | V | Initial Okhotsk | This study |
| HM2-R1 | Rib | 43.9 | 16.2 | -13.2 | 18.4 | 3.2 | R | – | Final Okhotsk | Ishida et al. (2000) |
| HM2-R2 | Rib | 45.0 | 16.5 | -13.0 | 18.6 | 3.2 | R | – | Middle Okhotsk | Ishida et al. (2000) |
| HM2-R3 | Rib | 45.0 | 16.3 | -13.7 | 18.7 | 3.2 | R | – | Final Okhotsk | Ishida et al. (2000) |
| HM2-HA-3 | Rib | 44.5 | 16.4 | -13.0 | 19.3 | 3.2 | HA | – | Middle Okhotsk | Uchida-Fukuhara et al. (2024) |
| HM2-HA-4 | Rib | 43.7 | 16.6 | -13.1 | 22.9 | 3.1 | HA | – | Middle Okhotsk | Ishida et al. (2000) |
| HN2-I-3 | Rib | 46.9 | 16.8 | -13.4 | 19.1 | 3.3 | I | – | Initial Okhotsk | Ishida et al. (2000) |

(Continued)

| ID | $^{14}C$ age | $^{14}C$ error | Calibrated $^{14}C$ - Aniva | | | Calibrated $^{14}C$ - Svobodnaya | | | Calibrated $^{14}C$ - Otaru | | |
|---|---|---|---|---|---|---|---|---|---|---|---|
| | | | Lower 2σ | Upper 2σ | Median | Lower 2σ | Upper 2σ | Median | Lower 2σ | Upper 2σ | Median |
| 93-1 | 1806 | 37 | 726 | 1046 | 886 | 771 | 1098 | 934.5 | 502 | 838 | 670 |
| HA2 NAT002 | 1689 | 20 | 880 | 1174 | 1027 | 907 | 1200 | 1053.5 | 650 | 940 | 795 |
| HA2 NAT003 | 1794 | 19 | 768 | 1052 | 910 | 785 | 1075 | 930 | 549 | 826 | 687.5 |
| HA2 NAT001 | 1995 | 20 | 583 | 847 | 715 | 615 | 878 | 746.5 | 325 | 641 | 483 |
| 2017HA0997 | 2271 | 28 | 261 | 576 | 418.5 | 305 | 609 | 457 | -25 | 328 | 151.5 |
| 2017HA0999 | 2262 | 28 | 266 | 584 | 425 | 318 | 628 | 473 | -14 | 343 | 164.5 |
| HA2 NAT004 | 2278 | 28 | 256 | 570 | 413 | 272 | 600 | 436 | -31 | 321 | 145 |
| HM2-R1 | 1721 | 37 | 826 | 1157 | 991.5 | 865 | 1188 | 1026.5 | 600 | 909 | 754.5 |
| HM2-R2 | 1797 | 37 | 732 | 1054 | 893 | 776 | 1096 | 936 | 519 | 845 | 682 |
| HM2-R3 | 1705 | 38 | 840 | 1171 | 1005.5 | 883 | 1203 | 1043 | 621 | 941 | 781 |
| HM2-HA-3 | 1777 | 37 | 771 | 1095 | 933 | 794 | 1120 | 957 | 552 | 865 | 708.5 |
| HM2-HA-4 | 1807 | 37 | 727 | 1044 | 885.5 | 765 | 1089 | 927 | 506 | 835 | 670.5 |
| HN2-I-3 | 2214 | 38 | 331 | 649 | 490 | 367 | 661 | 514 | 43 | 406 | 224.5 |

**Supplementary Table S2.** Stable isotopic data of dog skeletons from the Hamanaka 2 site.

| Lab ID | ID | %C | %N | $\delta^{13}C$ | $\delta^{15}N$ | C/N | Yield | Layer | Period | Name | Element | Grid |
|---|---|---|---|---|---|---|---|---|---|---|---|---|
| hd01 | 2011HA1751 | 42.4 | 15.5 | -12.6 | 17.4 | 3.2 | 11.2 | IVb | Initial Okhotsk | Dog | Mandible R | A03-d-3 |
| hd02 | 2011HA0858 | 40.4 | 14.6 | -13.4 | 16.8 | 3.2 | 6.2 | IV | Initial Okhotsk | Dog | Mandible R | B03-a-1 |
| hd03 | 2011HA2808-2811 | 42.9 | 15.7 | -12.7 | 16.8 | 3.2 | 10.4 | IIIb | Middle Okhotsk | Dog | Mandible R | C03-a-2 |
| hd04 | 2011HA1310 | 43.1 | 15.6 | -13.2 | 16.5 | 3.2 | 14.4 | IV | Initial Okhotsk | Dog | Mandible R | A03-d-1 |
| hd05 | 2011HA3930 | 36.3 | 13.1 | -13.7 | 16.3 | 3.2 | 4.0 | V | Initial Okhotsk | Dog | Mandible R | C03-c-1 |
| hd06 | 2011HA3384 | 38.4 | 13.9 | -12.9 | 17.2 | 3.2 | 6.4 | V | Initial Okhotsk | Dog | Mandible R | A03-c-4 |
| hd07 | 2011HA4906 | – | – | – | – | – | 0.5 | IVa | Initial Okhotsk | Dog | Mandible R | A03-ｂ-1 |
| hd08 | 2011HA1555 | 42.5 | 15.5 | -12.7 | 17.3 | 3.2 | 16.2 | IV | Initial Okhotsk | Dog | Mandible R | A03-a-3 |
| hd09 | 2011HA2825 | 43.5 | 15.8 | -13.1 | 16.7 | 3.2 | 12.8 | V | Initial Okhotsk | Dog | Mandible R | B03-d-4 |
| hd10 | 2011HA1210 | 42.4 | 15.3 | -13.0 | 17.2 | 3.2 | 9.4 | IV | Initial Okhotsk | Dog | Mandible R | A03-d-3 |
| hd11 | 2011HA0978 | 42.9 | 15.7 | -13.1 | 17.4 | 3.2 | 3.1 | IIIa | Middle Okhotsk | Dog | Mandible R | C03-b-4 |
| hd12 | 2011HA1560 | 43.4 | 15.6 | -13.0 | 16.9 | 3.3 | 6.0 | IVb | Initial Okhotsk | Dog | Mandible R | A03-d-3 |
| hd13 | 2011HA5011 | – | – | – | – | – | 0.1 | IVa | Initial Okhotsk | Dog | Mandible R | A03-b-4 |
| hd14 | 2011HA2737 | 44.1 | 16.1 | -13.4 | 16.4 | 3.2 | 12.8 | IIIb | Middle Okhotsk | Dog | Mandible R | C03-a-1 |
| hd15 | 2011HA5027 | 43.0 | 15.7 | -12.9 | 17.2 | 3.2 | 12.2 | IVb | Initial Okhotsk | Dog | Mandible R | A03-b-4 |
| hd16 | 2011HA1142 | 43.1 | 15.6 | -12.6 | 17.5 | 3.2 | 9.5 | IV | Initial Okhotsk | Dog | Mandible R | A03-b-2 |
| hd17 | 2011HA1317 | 43.3 | 15.7 | -12.8 | 16.9 | 3.2 | 10.6 | IVb | Initial Okhotsk | Dog | Mandible R | A03-d-3 |
| hd18 | 2011HA1608 | 44.2 | 16.1 | -13.2 | 16.4 | 3.2 | 12.0 | IVb | Initial Okhotsk | Dog | Mandible R | A03-d-1 |
| hd19 | 2014HA0310 | 45.2 | 16.4 | -13.6 | 17.3 | 3.2 | 14.7 | IIId | Middle Okhotsk | Dog | Mandible R | Z04-c-3 |
| hd20 | 2014HA0355 | 44.1 | 15.6 | -14.1 | 19.1 | 3.3 | 6.5 | IIId | Middle Okhotsk | Dog | Mandible R | Z03-b-2 |
| hd21 | 2014HA0436 | 45.9 | 16.8 | -13.4 | 19.3 | 3.2 | 11.6 | IIIc | Middle Okhotsk | Dog | Mandible R | Z03-b-3 |
| hd22 | 2014HA0625 | 45.6 | 16.5 | -13.5 | 18.3 | 3.2 | 8.9 | IIId | Middle Okhotsk | Dog | Mandible R | Z03-b-3 |
| hd23 | 2014HA1068 | 45.1 | 16.0 | -13.1 | 18.2 | 3.3 | 4.6 | VII | Epi-Jomon | Dog | Mandible R | A03-c-2 |

| | | | | | | | | | | | | |
|---|---|---|---|---|---|---|---|---|---|---|---|---|
| hd24 | 2014HA1139 | 44.3 | 16.1 | -12.8 | 16.8 | 3.2 | 6.4 | IIId | Middle Okhotsk | Dog | Mandible R | B03-a-2 |
| hd25 | 2014HA1194 | 45.2 | 16.0 | -13.8 | 18.4 | 3.3 | 5.9 | IIId | Middle Okhotsk | Dog | Mandible R | A04-d-2 |
| hd26 | 2014HA1202 | 45.1 | 16.5 | -13.5 | 18.7 | 3.2 | 10.5 | IIId | Middle Okhotsk | Dog | Mandible R | A04-c-1 |
| hd27 | 2014HA1279 | 43.5 | 15.2 | -13.2 | 17.4 | 3.3 | 3.2 | VII | Epi-Jomon | Dog | Mandible R | A03-a-3 |
| hd28 | 2014HA1487 | 45.1 | 16.6 | -13.5 | 16.3 | 3.2 | 9.7 | IIIb | Middle Okhotsk | Dog | Mandible R | A02-a-1 |
| hd29 | 2014HA1523 | 45.6 | 15.7 | -14.5 | 19.8 | 3.4 | 3.7 | IIIc | Middle Okhotsk | Dog | Mandible R | A02-c-1 |
| hd30 | 2014HA1762 | 44.9 | 15.5 | -13.7 | 18.1 | 3.4 | 2.8 | VIII | Epi-Jomon | Dog | Mandible R | A03-c-2 |
| hd31 | 2014HA1900 | 43.0 | 14.7 | -13.6 | 18.5 | 3.4 | 5.0 | IIIc | Middle Okhotsk | Dog | Mandible R | A04-c-1 |
| hd32 | 2014HA1969 | 45.1 | 15.8 | -14.1 | 19.5 | 3.3 | 6.1 | V | Initial Okhotsk | Dog | Mandible R | Z03-b-2 |
| hd33 | 2014HA1978 | 44.9 | 15.8 | -14.0 | 19.5 | 3.3 | 4.3 | IIIc | Middle Okhotsk | Dog | Mandible R | A04-c-1 |
| hd34 | 2014HA2086 | 46.6 | 16.7 | -13.2 | 16.8 | 3.3 | 3.4 | IV | Initial Okhotsk | Dog | Mandible R | A04-d-2 |
| hd35 | 2014HA2304 | 45.4 | 16.6 | -12.9 | 16.9 | 3.2 | 12.4 | IV | Initial Okhotsk | Dog | Mandible R | A04-a-3 |
| hd36 | 2014HA2420 | 46.0 | 16.4 | -13.0 | 19.3 | 3.3 | 7.5 | VIII | Epi-Jomon | Dog | Mandible R | A03-a-3 |
| hd37 | 2014HA2442 | 44.3 | 15.6 | -13.0 | 18.5 | 3.3 | 3.8 | VII | Epi-Jomon | Dog | Mandible R | B03-b-4 |
| hd38 | 2014HA2549 | 46.3 | 16.7 | -12.7 | 17.4 | 3.2 | 8.0 | IIIc | Middle Okhotsk | Dog | Mandible R | A02-b-4 |
| hd39 | 2014HA2613 | 45.1 | 15.9 | -13.4 | 16.4 | 3.3 | 5.4 | IIIc | Middle Okhotsk | Dog | Mandible R | A02-a-2 |
| hd40 | 2014HA2643 | 45.0 | 16.3 | -13.3 | 16.7 | 3.2 | 6.0 | IV | Initial Okhotsk | Dog | Mandible R | A04-d-2 |
| hd41 | 2014HA2752 | 44.8 | 16.0 | -12.8 | 16.6 | 3.3 | 6.4 | IIIc | Middle Okhotsk | Dog | Mandible R | A02-b-1 |
| hd42 | 2014HA2782 | 44.5 | 15.9 | -13.4 | 18.3 | 3.3 | 6.8 | IIIc | Middle Okhotsk | Dog | Mandible R | A02-b-1 |
| hd43 | 2011HA0645 | 44.1 | 15.9 | -13.2 | 20.1 | 3.2 | 14.5 | IV | Initial Okhotsk | Dog | Mandible R | B03-a-2 |
| hd44 | 2011HA0656 | 44.3 | 16.0 | -13.0 | 17.2 | 3.2 | 12.6 | III | Middle Okhotsk | Dog | Mandible R | A03-a-1 |
| hd45 | 2011HA0857 | 45.0 | 16.2 | -13.0 | 17.0 | 3.2 | 9.4 | IV | Initial Okhotsk | Dog | Mandible R | B03-a-1 |
| hd46 | 2011HA0859 | 42.1 | 15.2 | -13.1 | 17.2 | 3.2 | 17.7 | IV | Initial Okhotsk | Dog | Mandible R | B03-a-4 |
| hd47 | 2011HA1290 | 42.4 | 15.0 | -13.9 | 19.9 | 3.3 | 6.0 | IV | Initial Okhotsk | Dog | Mandible R | A03-b-2 |
| hd48 | 2011HA1378 | 41.8 | 15.0 | -13.2 | 18.4 | 3.2 | 12.4 | IV | Initial Okhotsk | Dog | Mandible R | A03-a-1 |

| | | | | | | | | | | | | |
|---|---|---|---|---|---|---|---|---|---|---|---|---|
| hd49 | 2011HA1591 | 42.0 | 15.4 | -13.3 | 18.6 | 3.2 | 11.9 | IVb | Initial Okhotsk | Dog | Mandible R | A03-d-2 |
| hd50 | 2011HA1952 | 42.4 | 15.4 | -13.0 | 18.0 | 3.2 | 13.9 | IV | Initial Okhotsk | Dog | Mandible R | A03-a-2 |
| hd51 | 2011HA2206 | 43.8 | 16.0 | -13.1 | 18.4 | 3.2 | 11.1 | IV | Initial Okhotsk | Dog | Mandible R | A03-a-2 |
| hd52 | 2011HA2217 | 42.0 | 15.4 | -13.3 | 19.0 | 3.2 | 14.9 | IV | Initial Okhotsk | Dog | Mandible R | A03-c-1 |
| hd53 | 2011HA2221 | 42.5 | 15.5 | -12.7 | 17.4 | 3.2 | 14.0 | IV | Initial Okhotsk | Dog | Mandible R | A03-c-1 |
| hd54 | 2011HA2424 | 43.1 | 15.5 | -13.4 | 17.6 | 3.2 | 8.9 | IVb | Initial Okhotsk | Dog | Mandible R | A03-c-1 |
| hd55 | 2011HA2604 | 41.7 | 15.2 | -13.1 | 18.0 | 3.2 | 9.0 | IVb | Initial Okhotsk | Dog | Mandible R | A03-c-4 |
| hd56 | 2011HA2620 | 41.9 | 15.3 | -12.6 | 17.3 | 3.2 | 11.7 | IV | Initial Okhotsk | Dog | Mandible R | A03-b-3 |
| hd57 | 2011HA2903 | 42.6 | 15.4 | -13.5 | 18.3 | 3.2 | 7.4 | V | Initial Okhotsk | Dog | Mandible R | B03-a-4 |
| hd58 | 2011HA2965 | 43.1 | 15.4 | -12.5 | 17.2 | 3.3 | 7.6 | V | Initial Okhotsk | Dog | Mandible R | B03-c-3 |
| hd59 | 2011HA3136 | 44.4 | 16.1 | -13.3 | 18.3 | 3.2 | 10.6 | V | Initial Okhotsk | Dog | Mandible R | A03-d-3 |
| hd60 | 2011HA3147 | 42.1 | 15.0 | -13.4 | 19.5 | 3.3 | 6.3 | V | Initial Okhotsk | Dog | Mandible R | A03-c-1 |
| hd61 | 2011HA3225 | 41.9 | 15.1 | -13.2 | 18.7 | 3.2 | 7.9 | V | Initial Okhotsk | Dog | Mandible R | B03-c-2 |
| hd62 | 2011HA3915 | 42.6 | 15.3 | -13.3 | 19.5 | 3.3 | 9.4 | V | Initial Okhotsk | Dog | Mandible R | A03-a-3 |
| hd63 | 2011HA4152 | 43.8 | 15.5 | -12.7 | 18.5 | 3.3 | 11.3 | V | Initial Okhotsk | Dog | Mandible R | C03-b-1 |
| hd64 | 2011HA4179 | 41.4 | 14.8 | -13.7 | 17.1 | 3.3 | 5.4 | V | Initial Okhotsk | Dog | Mandible R | A03-d-3 |
| hd65 | 2011HA4911 | 43.0 | 15.7 | -13.0 | 17.5 | 3.2 | 15.1 | IVa | Initial Okhotsk | Dog | Mandible R | A03-b-1 |
| hd66 | 2011HA4961 | 42.1 | 15.3 | -14.3 | 17.7 | 3.2 | 9.6 | V | Initial Okhotsk | Dog | Mandible R | A03-b-1 |
| hd67 | 2011HA5029 | 43.5 | 15.8 | -12.8 | 17.9 | 3.2 | 11.2 | IVb | Initial Okhotsk | Dog | Mandible R | A03-b-4 |
| hd68 | 2011DG1 | 42.4 | 15.3 | -12.5 | 17.8 | 3.2 | 6.7 | VIII | Epi-Jomon | Dog | Mandible R | Test pit |
| hd69 | 2011DG2 | 42.9 | 15.7 | -12.9 | 17.4 | 3.2 | 12.8 | IV | Initial Okhotsk | Dog | Mandible R | Test pit |
| hd70 | 2011DG3 | 41.8 | 15.3 | -13.5 | 18.5 | 3.2 | 7.0 | IV | Initial Okhotsk | Dog | Mandible R | B03-a-4 |
| hd71 | 2011DG15 | 41.8 | 15.4 | -13.3 | 17.8 | 3.2 | 11.1 | IVb | Initial Okhotsk | Dog | Mandible R | A03-c-4 |
| hd72 | 2011DG16 | 40.9 | 14.7 | -13.0 | 17.6 | 3.2 | 7.9 | IVb | Initial Okhotsk | Dog | Mandible R | A03-d-2 |
| hd73 | 2011DG17 | 42.1 | 15.1 | -12.7 | 16.5 | 3.2 | 9.4 | II | Final Okhotsk | Dog | Mandible R | A03-d-2 |

| | | | | | | | | | | | | |
|---|---|---|---|---|---|---|---|---|---|---|---|---|
| hd74 | 2013HA0484 | 42.5 | 15.4 | -12.4 | 17.6 | 3.2 | 11.4 | VII | Epi-Jomon | Dog | Mandible R | A03-d-2 |
| hd75 | 2013HA0905 | 41.7 | 15.2 | -13.2 | 18.8 | 3.2 | 9.0 | IIIb | Middle Okhotsk | Dog | Mandible R | A04-c-3 |
| hd76 | 2013HA0908 | 41.9 | 15.2 | -12.9 | 18.7 | 3.2 | 10.3 | IIIb | Middle Okhotsk | Dog | Mandible R | A04-c-4 |
| hd77 | 2013HA0951 | 42.6 | 15.5 | -12.9 | 18.3 | 3.2 | 14.6 | IIIb | Middle Okhotsk | Dog | Mandible R | A04-c-4 |
| hd78 | 2013HA2062 | 42.1 | 15.0 | -13.7 | 18.5 | 3.3 | 8.9 | IIIb | Middle Okhotsk | Dog | Mandible R | A04-d-4 |
| hd79 | 2013HA2063 | 42.2 | 15.2 | -13.4 | 18.8 | 3.2 | 11.5 | IIIb | Middle Okhotsk | Dog | Mandible R | A04-d-4 |
| hd80 | 2013HA2082 | 42.9 | 15.6 | -12.9 | 16.9 | 3.2 | 12.8 | IIIb | Middle Okhotsk | Dog | Mandible R | A04-d-4 |
| J14 | 2013HA0014 | 34.5 | 12.1 | -12.8 | 16.3 | 3.3 | 10.2 | III | Middle Okhotsk | Dog | Vertebra | A03-c1 |
| J16 | 2013HA0016 | 38.9 | 13.3 | -15.1 | 16.1 | 3.4 | 7.5 | II | Final Okhotsk | Dog | Humerus/Femur | A04-c |
| J18 | 2013HA0018 | 39.1 | 14.0 | -13.8 | 19.3 | 3.3 | 10.6 | IIIb | Middle Okhotsk | Dog | Rib | A04-c4 |
| J20 | 2013HA0020 | 42.1 | 15.0 | -13.8 | 17.5 | 3.3 | 9.8 | IIIc | Middle Okhotsk | Dog | Vertebra | A04-c3 |
| J22 | 2013HA0022 | 27.9 | 9.7 | -13.1 | 17.4 | 3.4 | 12.6 | VII | Epi-Jomon | Dog | Rib | A03-d2 |

**Supplementary Table S3.** Stable isotopic data of pig skeletons from the Hamanaka 2 site.

| Lab ID | ID | %C | %N | $\delta^{13}C$ | $\delta^{15}N$ | C/N | Yield | Layer | Period | Name | Element | Grid |
|---|---|---|---|---|---|---|---|---|---|---|---|---|
| hp01 | 2011HA0753 | 43.6 | 15.9 | -18.5 | 10.9 | 3.2 | 7.8 | – | – | Pig | Mandible L | C03-c-4 |
| hp02 | 2011HA1594 | 43.4 | 15.6 | -17.9 | 12.7 | 3.2 | 6.9 | IV | Initial Okhotsk | Pig | Mandible L | B03-d-3 |
| hp03 | 2011HA1812 | 43.0 | 15.7 | -16.5 | 14.0 | 3.2 | 12.4 | IIIb | Middle Okhotsk | Pig | Mandible L | C03-d-4 |
| hp04 | 2011HA1052 | 42.3 | 15.3 | -16.8 | 14.6 | 3.2 | 8.5 | IV | Initial Okhotsk | Pig | Mandible L | B03-d-1 |
| hp05 | 2011HA1879 | 43.9 | 15.8 | -14.9 | 15.7 | 3.2 | 8.7 | IIIb | Middle Okhotsk | Pig | Mandible L | B03-b-1 |
| hp06 | 2014HA0381 | 46.2 | 16.6 | -19.6 | 14.0 | 3.3 | 6.8 | IIIc | Middle Okhotsk | Pig | Mandible L | B03-a-3 |
| hp07 | 2014HA1568 | 46.3 | 16.9 | -14.9 | 15.7 | 3.2 | 11.9 | IIIb | Middle Okhotsk | Pig | Mandible L | A02-a-1 |
| hp08 | 2014HA2153 | 49.9 | 17.8 | -13.6 | 17.2 | 3.3 | 9.6 | IIIb | Middle Okhotsk | Pig | Mandible L | A02-b-1 |
| hp09 | 2014HA2226 | 43.4 | 15.8 | -19.6 | 10.3 | 3.2 | 13.0 | IIId | Middle Okhotsk | Pig | Mandible L | A04-d-2 |
| hp10 | 2011HA0732 | 45.4 | 16.3 | -18.1 | 13.2 | 3.2 | 10.7 | III | Middle Okhotsk | Pig | Mandible L | B03-d-3 |
| hp11 | 2011PG8 | 43.5 | 15.7 | -16.0 | 15.4 | 3.2 | 10.1 | IV | Initial Okhotsk | Pig | Mandible L | A03-a-4 |
| hp12 | 2011PG9 | 41.3 | 14.9 | -18.2 | 10.9 | 3.2 | 8.4 | IVb | Initial Okhotsk | Pig | Mandible L | A03-d-3 |
| hp13 | 2014HA0646 | 44.8 | 16.2 | -15.5 | 15.7 | 3.2 | 10.0 | IIIb | Middle Okhotsk | Pig | Mandible R | A02-b-1 |
| hp14 | 2011General | 44.9 | 16.2 | -13.3 | 16.4 | 3.2 | 9.9 | IIIa | Middle Okhotsk | Pig | Vertebra | C03-c-1 |
| hp15 | 2011HA2529 | 45.3 | 16.3 | -17.2 | 12.1 | 3.2 | 9.4 | IVb | Initial Okhotsk | Pig | Mandible R | A03-c-4 |
| hp16 | 2011HA1727 | 45.5 | 16.6 | -14.5 | 16.0 | 3.2 | 13.0 | IV | Initial Okhotsk | Pig | Mandible R | A03-a-3 |
| hp17 | 2011HA1968 | 44.9 | 16.2 | -17.7 | 13.4 | 3.2 | 13.3 | IV | Initial Okhotsk | Pig | Mandible R | A03-c-1 |
| hp18 | 2013HA1909 | 43.5 | 15.5 | -14.9 | 17.7 | 3.3 | 6.2 | IIIb | Middle Okhotsk | Pig | Mandible R | A04-c-4 |
| hp19 | 2014HA0195&0194 | 45.5 | 16.3 | -17.4 | 13.8 | 3.3 | 9.3 | IIIb | Middle Okhotsk | Pig | Mandible R | Z03-b-3 |
| hp20 | 2011G1 | 45.1 | 16.2 | -17.0 | 13.3 | 3.3 | 6.2 | IV | Initial Okhotsk | Pig | Mandible R | Test pit |
| hp21 | 2011G2 | 45.9 | 16.5 | -14.4 | 16.1 | 3.3 | 7.0 | III | Middle Okhotsk | Pig | Mandible R | C03-d-2 |
| hp22 | 2011HA3861 | 45.4 | 16.4 | -21.1 | 6.9 | 3.2 | 12.8 | V | Initial Okhotsk | Pig | Maxilla R | A03-b-3 |
| hp23 | 2011HA2474 | 44.6 | 15.3 | -12.8 | 18.2 | 3.4 | 14.9 | IIIa | Middle Okhotsk | Pig | Maxilla L | C03-b-1 |

| | | | | | | | | | | | | |
|---|---|---|---|---|---|---|---|---|---|---|---|---|
| hp24 | 2011HA1279 | 46.0 | 16.3 | -19.2 | 10.3 | 3.3 | 8.2 | IIIa | Middle Okhotsk | Pig | Maxilla L | C03-b-3 |
| hp25 | 2015HA0268 | 44.4 | 15.9 | -16.7 | 14.0 | 3.3 | 8.5 | IIId | Middle Okhotsk | Pig | Radius R | A02-b-3 |
| hp26 | 2011Humerus６ | 47.0 | 17.2 | -18.3 | 10.2 | 3.2 | 11.7 | V | Initial Okhotsk | Pig | Humerous | Test pit |
| hp27 | 2011HA4740 | 43.4 | 15.7 | -18.6 | 12.0 | 3.2 | 7.6 | V | Initial Okhotsk | Pig | Lower vertebra | A03-d-3 |
| hp28 | 2011General | 44.7 | 16.1 | -14.4 | 16.5 | 3.2 | 11.3 | IIIa | Middle Okhotsk | Pig | Rib | C03-b-3 |
| hp29 | 2011General | 45.0 | 16.3 | -14.3 | 15.8 | 3.2 | 11.5 | IIIa | Middle Okhotsk | Pig | Vertebra | C03-c-1 |
| hp30 | 2015HASG0491 | 45.1 | 16.2 | -18.5 | 13.5 | 3.3 | 8.0 | IIc | Final Okhotsk | Pig | Tibia | B02-c-1 |
| hp31 | 2011HA1771 | 43.8 | 15.6 | -15.3 | 15.6 | 3.3 | 9.4 | IIIb | Middle Okhotsk | Pig | Cranium | C03-d-1 |
| hp32 | 2011HA0896＆HA0814 | 42.4 | 15.3 | -17.1 | 14.2 | 3.2 | 10.3 | IIIa | Middle Okhotsk | Pig | Cranium | C03-c-4/b-4 |
| hp33 | 2011HA1893 | 41.2 | 14.9 | -14.4 | 15.0 | 3.2 | 7.4 | IV | Initial Okhotsk | Pig | Mandible | A03-a-2 |

**Supplementary Table S4.** Stable isotopic data of faunal skeletons, other than domesticated dogs and pigs, from the Hamanaka 2 site.

| Lab ID | ID | %C | %N | $\delta^{13}C$ | $\delta^{15}N$ | C/N | Yield | Layer | Period | Name | Element | Grid |
|---|---|---|---|---|---|---|---|---|---|---|---|---|
| ha2f31 | 2011HA2780 | 45.3 | 16.4 | -13.8 | 14.3 | 3.2 | 6.4 | IIIa/b | Middle Okhotsk | Whale | Maxilla | C03-a1 |
| ha2f32 | 2011HA2603 | 40.9 | 14.9 | -12.4 | 17.9 | 3.2 | 7.3 | IIIb | Middle Okhotsk | Sea lion | Maxilla | C03-b1 |
| ha2f33 | 2011HA0732 | 41.6 | 15.2 | -13.7 | 18.3 | 3.2 | 14.5 | III | Middle Okhotsk | Sea lion | Calcaneus | B03-d3 |
| ha2f34 | 2013HA1986 | 41.5 | 14.4 | -13.9 | 18.8 | 3.4 | 3.6 | IIIb | Middle Okhotsk | Steller sea lion | Ulna | A02-b4 |
| ha2f35 | 2013HA1052 | 41.3 | 15.0 | -14.2 | 18.3 | 3.2 | 11.9 | IIIc | Middle Okhotsk | Fur seal | Axis | Z04-c3 |
| ha2f36 | 2013HA1867 | 39.7 | 14.4 | -14.3 | 17.4 | 3.2 | 6.7 | IIIb | Middle Okhotsk | Steller sea lion | Femur | A02-d2 |
| ha2f37 | 2013HA1377 | 41.6 | 15.1 | -13.5 | 16.6 | 3.2 | 11.0 | IIIc | Middle Okhotsk | Spotted seal | Humerus | A03-c3 |
| ha2f38 | 2011HA0295 | 42.2 | 15.4 | -14.8 | 16.9 | 3.2 | 11.9 | III | Middle Okhotsk | Fur seal | Ulna | A03-a3 |
| ha2f39 | 2013HA1673 | 42.3 | 15.2 | -13.2 | 16.6 | 3.2 | 15.6 | IIIb | Middle Okhotsk | Fox | Mandible | Z03-b2 |
| ha2f40 | 2013HA0962 | 41.7 | 15.0 | -17.7 | 11.3 | 3.2 | 12.3 | IIIa | Middle Okhotsk | Marten | Mandible | A04-d3 |
| ha2f41 | 2015HA0923 | 45.8 | 16.8 | -13.7 | 18.0 | 3.2 | 11.4 | IIIa | Middle Okhotsk | Steller sea lion | Scapula | Z03-c2 |
| ha2f42 | 2015HASG470 | 44.1 | 15.6 | -16.8 | 12.5 | 3.3 | 5.0 | VIII | Epi-Jomon | Fox | – | A03-a1 |
| ha2f43 | 2015HASG409 | 44.4 | 16.0 | -17.0 | 12.9 | 3.2 | 5.2 | VIII | Epi-Jomon | Fox | – | A03-a1 |
| ha2f44 | 2015HASG520 | 44.0 | 15.6 | -13.4 | 17.2 | 3.3 | 5.7 | IIc | Final Okhotsk | Fox | – | B02-a3 |
| ha2f45 | 2015HASG508 | 44.2 | 16.1 | -15.5 | 11.2 | 3.2 | 9.4 | VIII | Epi-Jomon | Fox | – | A03-a2 |
| ha2f46 | 2015HASG528 | – | – | – | – | – | 0.0 | VIII | Epi-Jomon | Fox | – | A03-a4 |
| ha2f47 | 2015HASG079 | 43.8 | 15.6 | -15.5 | 13.4 | 3.3 | 7.7 | V | Initial Okhotsk | Whale | – | A02-b4 |
| ha2f48 | 2015HASG397 | 43.5 | 15.6 | -13.9 | 13.3 | 3.3 | 7.2 | IIa | Final Okhotsk | Fox | – | B02-a1 |
| ha2f49 | 2015HASG499 | 43.6 | 15.6 | -16.3 | 13.1 | 3.3 | 8.4 | VIII | Epi-Jomon | Fox | – | A03-a2 |
| ha2f50 | 2015HASG517 | 44.2 | 16.0 | -13.2 | 16.2 | 3.2 | 10.7 | IIc | Final Okhotsk | Fox | – | B02-b4 |
| ha2f51 | 2015HASG469 | 44.4 | 16.0 | -14.5 | 14.5 | 3.2 | 8.6 | VIII | Epi-Jomon | Fox | – | A03-a1 |
| ha2f52 | 2013HA0025 | 43.0 | 15.8 | -14.1 | 14.8 | 3.2 | 6.1 | I | Ainu | Dolphin | – | A04-d3 |
| ha2f53 | 2013HA0024 | 43.6 | 15.1 | -14.6 | 14.9 | 3.4 | 4.9 | IIa | Final Okhotsk | Dolphin | – | A02-d |

| | | | | | | | | | | | | |
|---|---|---|---|---|---|---|---|---|---|---|---|---|
| ha2f54 | 2013HA0023 | 43.5 | 15.1 | -15.8 | 15.9 | 3.4 | 6.0 | IIIa | Middle Okhotsk | Dolphin | – | Z04-c3 |
| ha2f55 | 2013HA0035 | 43.7 | 15.5 | -13.8 | 17.9 | 3.3 | 7.1 | IIc | Final Okhotsk | Earless seal | – | A02-a |
| ha2f56 | 2015HASG524 | 43.0 | 15.2 | -14.5 | 17.3 | 3.3 | 5.8 | VIII | Epi-Jomon | Fur seal | – | A03-a4 |
| ha2f57 | 2015HASG441 | 44.0 | 16.0 | -14.5 | 15.1 | 3.2 | 10.5 | IIIb | Middle Okhotsk | Ribbon_seal | – | Z03-c2 |
| ha2f58 | 2015HASG512 | 44.8 | 16.1 | -14.8 | 17.8 | 3.2 | 6.6 | IIIb | Middle Okhotsk | Fur seal | – | Z03-c2 |
| ha2f59 | – | 41.6 | 15.0 | -14.4 | 16.8 | 3.2 | 7.2 | IIIc | Middle Okhotsk | Red throated loon | – | A02-b4 |
| ha2f60 | – | 42.0 | 15.2 | -16.5 | 20.8 | 3.2 | 11.3 | IIIb | Middle Okhotsk | Seagull | – | Z03-b2 |
| ha2f61 | 2013HA1871 | 41.4 | 14.8 | -14.0 | 13.9 | 3.3 | 7.2 | IIIb | Middle Okhotsk | Eagle | – | A02-c1 |
| ha2f62 | – | 42.3 | 14.9 | -14.7 | 16.0 | 3.3 | 14.6 | IIIa | Middle Okhotsk | Goose | – | A04-d2 |
| ha2f63 | – | 39.8 | 14.4 | -15.9 | 11.0 | 3.2 | 7.1 | III | Middle Okhotsk | Crow | – | Z02-b2 |
| ha2f64 | 2015HA879 | 39.3 | 14.1 | -12.8 | 16.7 | 3.3 | 6.6 | VIII | Epi-Jomon | Crow | – | C03-a3 |
| ha2f65 | – | 40.2 | 14.5 | -15.3 | 16.5 | 3.2 | 8.2 | IIIc | Middle Okhotsk | Guillemot | – | A02-b4 |
| ha2f66 | – | 40.5 | 14.6 | -15.4 | 16.3 | 3.2 | 6.6 | VIII | Epi-Jomon | Guillemot | – | A03-b3 |
| ha2f67 | 2013HA752 | 39.9 | 14.3 | -13.5 | 15.2 | 3.2 | 6.0 | IIIa | Middle Okhotsk | Cormorant | – | A04-d2 |
| ha2f68 | 2013HA2142/2162 | 41.1 | 14.9 | -14.2 | 14.9 | 3.2 | 7.4 | VIII | Epi-Jomon | Cormorant | – | A03-d2/A03-c1 |
| ha2f69 | – | 41.1 | 14.8 | -14.1 | 18.3 | 3.2 | 5.6 | VIII | Epi-Jomon | Albatros | – | B03-a3/a4 |
| ha2f70 | 2013HA1842 | 40.8 | 14.8 | -13.4 | 16.7 | 3.2 | 5.7 | IIIb | Middle Okhotsk | Albatros | – | A02-d2 |
| J53 | 2013HA0053 | 35.1 | 12.6 | -13.3 | 19.4 | 3.3 | 14.6 | I | Ainu | Sea lion | – | A04-d3 |
| J55 | 2013HA0055 | 35.9 | 12.7 | -14.3 | 18.2 | 3.3 | 9.6 | – | – | Sea lion | Rib | A02-b |
| J61 | 2013HA0061 | 39.8 | 13.9 | -14.6 | 18.8 | 3.3 | 11.7 | I | Ainu | Sea lion | Vertebrae | A02-c |
| J63 | 2013HA0063 | 30.4 | 10.8 | -14.2 | 17.8 | 3.3 | 9.4 | IIc | Final Okhotsk | Sea lion | Rib | A04-d4 |
| J65 | 2013HA0065 | 39.6 | 14.0 | -14.3 | 18.2 | 3.3 | 17.7 | I | Ainu | Sea lion | Rib | A02-d |
| J67 | 2013HA0067 | 36.8 | 13.0 | -14.3 | 18.0 | 3.3 | 14.0 | I | Ainu | Sea lion | Rib | A04-c |
| J68 | 2013HA0068 | 30.2 | 10.1 | -14.5 | 18.8 | 3.5 | 9.5 | I | Ainu | Sea lion | Ulna | A04-c3 |
| J69 | 2013HA0069 | 39.9 | 13.7 | -14.9 | 18.4 | 3.4 | 6.5 | I | Ainu | Sea lion | Phalanx | A02-c |

| | | | | | | | | | | | | |
|---|---|---|---|---|---|---|---|---|---|---|---|---|
| J71 | 2013HA0071 | 38.3 | 13.3 | -15.7 | 19.4 | 3.4 | 7.1 | IIa | Final Okhotsk | Sea lion | – | A02-d2 |
| J37 | 2013HA0037 | 25.4 | 8.4 | -15.9 | 19.4 | 3.5 | 8.6 | I | Ainu | Spotted seal | Femur | A02-c |
| J38 | 2013HA0038 | 29.9 | 10.2 | -16.0 | 19.6 | 3.4 | 10.0 | I | Ainu | Spotted seal | Humerus | A04-c4 |
| J40 | 2013HA0040 | 43.1 | 15.2 | -15.3 | 20.2 | 3.3 | 8.9 | IIa | Final Okhotsk | Spotted seal | Vertebrae | A03-c |
| J41 | 2013HA0041 | 34.0 | 12.2 | -14.1 | 17.7 | 3.3 | 10.3 | IIIa | Middle Okhotsk | Spotted seal | Metapoidial | Z-04-c3 |
| J43 | 2013HA0043 | 44.2 | 15.5 | -15.5 | 20.2 | 3.3 | 7.6 | IIa | Final Okhotsk | Spotted seal | Vertebrae | A02-c |
| J44 | 2013HA0044 | 41.6 | 14.7 | -14.8 | 18.4 | 3.3 | 4.8 | IIa | Final Okhotsk | Spotted seal | Vertebrae | A02-b |
| J45 | 2013HA0045 | 39.2 | 13.7 | -15.2 | 19.2 | 3.3 | 9.1 | IIa | Final Okhotsk | Spotted seal | Rib | A03-c3 |
| J46 | 2013HA0046 | 33.9 | 11.7 | -13.9 | 17.3 | 3.4 | 9.0 | IIa | Final Okhotsk | Spotted seal | Rib | A02-d |
| J49 | 2013HA0049 | 33.8 | 12.0 | -13.3 | 17.1 | 3.3 | 16.5 | I | Ainu | Spotted seal | Ulna | A02-c |
| J50 | 2013HA0050 | 38.9 | 13.8 | -14.3 | 16.0 | 3.3 | 8.5 | IIIa | Middle Okhotsk | Spotted seal | Femur | A04-d2 |
| J51 | 2013HA0051 | 31.1 | 10.8 | -13.2 | 17.3 | 3.4 | 9.3 | I | Ainu | Spotted seal | Humerus | A04-c3 |
| J88 | 2013HA0088 | 37.5 | 13.3 | -16.2 | 11.7 | 3.3 | 12.5 | II | Final Okhotsk | Whale | – | A04-c |
| J06 | 2013HA0006 | 39.3 | 13.9 | -13.9 | 17.7 | 3.3 | 15.6 | IIc | Final Okhotsk | Albatros | Humerus | A02-a |
| J80 | 2013HA0080 | 38.7 | 13.5 | -16.2 | 16.3 | 3.3 | 10.1 | IIa | Final Okhotsk | Puffin | – | A03-c3 |

**Supplementary Table S5.** Stable isotopic data of modern food samples from Rebun Island. Raw values without correction for the Suess effect were shown for $\delta^{13}C$ values.

| Lab ID | %C | %N | $\delta^{13}C$ | $\delta^{15}N$ | C/N | Period | Classification | Name | Scientific name |
|---|---|---|---|---|---|---|---|---|---|
| RF02 | 51.7 | 0.8 | -27.5 | -0.2 | 74.7 | Modern | C3 | Crowberry | *Empetrum nigrum var. japonicum* |
| RF04 | 41.4 | 0.4 | -26.6 | 1.5 | 119.0 | Modern | C3 | Rosehip | *Rosa rugosa* |
| RF24 | 51.7 | 0.7 | -27.0 | 0.6 | 85.6 | Modern | C3 | Crowberry | *Empetrum nigrum var. japonicum* |
| RF25 | 49.0 | 0.3 | -28.1 | 1.2 | 225.1 | Modern | C3 | Crowberry | *Empetrum nigrum var. japonicum* |
| RF27 | 41.9 | 0.7 | -26.5 | 1.2 | 72.4 | Modern | C3 | Rosehip | *Rosa rugosa* |
| RF32 | 41.6 | 0.9 | -25.9 | 1.4 | 56.8 | Modern | C3 | Rosehip | *Rosa rugosa* |
| RF36 | 36.3 | 0.8 | -25.5 | 1.0 | 49.8 | Modern | C3 | Grapevine | *Vitis coignetiae* |
| RF37 | 38.1 | 0.8 | -26.1 | -1.1 | 52.5 | Modern | C3 | Grapevine | *Vitis coignetiae* |
| RF38 | 38.3 | 1.0 | -29.1 | 1.3 | 44.0 | Modern | C3 | Grapevine | *Vitis coignetiae* |
| RF41 | 45.0 | 2.0 | -29.7 | 6.0 | 26.0 | Modern | C3 | Kiwi berry | *Actinidia arguta* |
| RF42 | 41.4 | 1.4 | -26.6 | 6.0 | 33.9 | Modern | C3 | Kiwi berry | *Actinidia arguta* |
| RF43 | 45.0 | 1.8 | -31.3 | 2.1 | 29.8 | Modern | C3 | Kiwi berry | *Actinidia arguta* |
| RF01 | 41.6 | 10.0 | -21.1 | 6.3 | 4.8 | Modern | MS | Northern sea urchin | *Strongylocentrotus nudus* |
| RF09 | 39.1 | 9.4 | -19.7 | 6.1 | 4.9 | Modern | MS | Bafun sea urchin | *Hemicentrotus pulcherrimus* |
| RF10 | 40.7 | 10.0 | -18.1 | 5.8 | 4.7 | Modern | MS | Bafun sea urchin | *Hemicentrotus pulcherrimus* |
| RF17 | 41.7 | 9.6 | -19.5 | 4.6 | 5.1 | Modern | MS | Northern sea urchin | *Strongylocentrotus nudus* |
| RF18 | 43.5 | 10.0 | -19.8 | 5.2 | 5.1 | Modern | MS | Northern sea urchin | *Strongylocentrotus nudus* |
| RF20 | 43.0 | 10.6 | -18.8 | 4.9 | 4.7 | Modern | MS | Bafun sea urchin | *Hemicentrotus pulcherrimus* |
| RF23 | 40.7 | 12.7 | -18.2 | 6.6 | 3.7 | Modern | MS | Abalone | *Haliotis sp.* |

**Supplementary Table S6.** Result of SIAR calculation for the proportions of protein contributions from each food source in different animals from the Hamanaka 2 site. For the Pig: Total, all pig individuals; Higher and Lower, each 7 pig individuals with the highest or lowest $\delta^{15}N$ values, respectively; Initial and Middle, pigs fromthe the Initial or Middle Okhotsk periods, respectively. For the Fox: Total, all fox individuals; Epi-Jomon and Okhotsk & Ainu, foxes from Epi-Jomon or Okhotsk/Ainu periods, respectively.

| | | C3 plants | | | Marine fish | | | Marine mammal/Dogs | | |
|---|---|---|---|---|---|---|---|---|---|---|
| | | Median | Lower | Upper | Median | Lower | Upper | Median | Lower | Upper |
| Pig | Total | 41.5 | 36.4 | 46.0 | 1.5 | 1.2 | 35.2 | 51.5 | 26.8 | 55.9 |
| | Lower | 64.5 | 56.6 | 71.9 | 1.5 | 1.0 | 34.4 | 28.5 | 4.9 | 34.7 |
| | Higher | 16.5 | 11.5 | 21.7 | 41.5 | 22.7 | 63.0 | 41.5 | 22.8 | 58.2 |
| | Initial | 49.5 | 40.7 | 56.1 | 1.5 | 1.4 | 44.8 | 40.5 | 9.7 | 48.7 |
| | Middle | 33.5 | 27.3 | 39.4 | 30.5 | 2.7 | 47.1 | 39.5 | 23.2 | 60.4 |
| Dog | | 8.5 | 7.1 | 9.7 | 52.5 | 47.0 | 58.7 | 39.5 | 33.5 | 44.1 |
| Human | | 5.5 | 1.8 | 9.6 | 45.5 | 26.2 | 57.6 | 50.5 | 38.0 | 66.8 |
| Fox | Total | 30.5 | 23.4 | 38.8 | 36.5 | 15.3 | 68.6 | 33.5 | 4.3 | 49.7 |
| | Epi-Jomon | 42.5 | 35.1 | 48.5 | 33.5 | 7.0 | 54.4 | 23.5 | 6.6 | 47.5 |
| | Okhotsk & Ainu | 14.5 | 5.0 | 25.2 | 43.5 | 24.2 | 81.0 | 38.5 | 6.3 | 59.8 |

**Supplementary Table S7.** Summary of stable isotopic results of human skeletons from the Hamanaka 2 site.

| | | $\delta^{13}C$ | | | | $\delta^{15}N$ | | | | n |
|---|---|---|---|---|---|---|---|---|---|---|
| | | Mean | SD | Minimum | Maximum | Mean | SD | Minimum | Maximum | |
| Adult | Total | -13.2 | 0.2 | -13.7 | -12.9 | 19 | 0.5 | 18.3 | 19.7 | 9 |
| | Female | -13.1 | 0.2 | -13.4 | -12.9 | 18.9 | 0.4 | 18.3 | 19.3 | 5 |
| | Male | -13.2 | 0.1 | -13.2 | -13.1 | 19 | 0.9 | 18.4 | 19.7 | 2 |
| | Unknown sex | -13.5 | 0.3 | -13.7 | -13.3 | 19.2 | 0.2 | 19.1 | 19.4 | 2 |
| Non-adult | | -13.7 | 0.5 | -14.4 | -13.1 | 20 | 1.9 | 18.7 | 22.9 | 4 |

**Supplementary Table S8.** Stable isotopic data of the tooth enamel increments of a pig individual (2011HA1894) from the Hamanaka 2 site. The distance from the cervical edge of $I_1$ is also shown.

| Tooth | Number | Distance (mm) | $\delta^{13}C$ | $\delta^{18}O$ |
|---|---|---|---|---|
| $I_1$ | 1 | 40.72 | -8.7 | -8.2 |
| | 2 | 31.28 | -8.4 | -6.8 |
| | 3 | 29.76 | -8.4 | -6.1 |
| | 4 | 28.01 | -8.8 | -6.1 |
| | 5 | 26.46 | -9.1 | -5.9 |
| | 6 | 25.03 | – | – |
| | 7 | 23.17 | – | – |
| | 8 | 21.52 | -9.8 | -5.5 |
| | 9 | 20.17 | -10.1 | -5.5 |
| | 10 | 18.53 | -10.3 | -5.4 |
| | 11 | 17.14 | -10.7 | -5.3 |
| | 12 | 15.64 | -10.6 | -5.1 |
| | 13 | 14.25 | -10.8 | -5.0 |
| | 14 | 12.42 | -10.9 | -4.6 |
| | 15 | 10.69 | -11.1 | -4.6 |
| | 16 | 9.51 | -11.4 | -4.6 |
| | 17 | 7.40 | -10.7 | -4.5 |
| | 18 | 5.87 | -9.9 | -4.5 |
| $I_2$ | 1 | 45.42 | -9.3 | -6.3 |
| | 2 | 43.83 | -9.5 | -5.9 |
| | 3 | 41.93 | -9.7 | -5.7 |

| | | | |
|---|---|---|---|
| 4 | 40.12 | -10.1 | -5.5 |
| 5 | 38.37 | -10.2 | -5.4 |
| 6 | 36.56 | -10.5 | -5.6 |
| 7 | 35.19 | -10.5 | -5.6 |
| 8 | 33.47 | -10.5 | -5.7 |
| 9 | 31.53 | -10.4 | -5.6 |
| 10 | 29.73 | -10.3 | -5.9 |
| 11 | 27.74 | -10.2 | -6.1 |
| 12 | 25.77 | -10.0 | -6.3 |
| 13 | 23.97 | -10.0 | -6.8 |
| 14 | 22.06 | -9.9 | -6.9 |
| 15 | 20.51 | -9.8 | -7.4 |
| 16 | 18.26 | -9.7 | -7.7 |
| 17 | 15.98 | -8.9 | -7.9 |
| 18 | 13.73 | -8.8 | -8.5 |
| 19 | 10.88 | -7.8 | -8.8 |
| 20 | 8.69 | -7.0 | -9.0 |
